# Hierarchical latent representations reveal protein organization for functional discovery and design

**DOI:** 10.64898/2026.02.14.705947

**Authors:** Zhengyang Guo, Zi Wang, Shimin Wang, Yongping Chai, Kaiming Xu, Ming Li, Wei Li, Guangshuo Ou

## Abstract

Proteins can preserve conserved functions despite extensive sequence and structural divergence, suggesting that functional organization is governed by distributed constraints not captured by conventional representations. Here we develop a hierarchical sequence-based representation framework that compresses proteins into context-dependent latent states while preserving multiscale organizational information. Using this framework, we identified previously uncharacterized ciliary proteins lacking detectable sequence and structure homology, including ADMAP1, which is required for normal sperm axonemal organization and motility in mice. Discrete latent protein states captured species-level organizational signatures correlated with major evolutionary groups and revealed expansion of intrinsically disordered regulatory environments in eukaryotes. Autoregressive sampling within this latent space further enabled design of synthetic actin-remodeling proteins that maintained robust F-actin severing activity despite extensive sequence rewiring across key functional interfaces. These findings demonstrate that distributed protein organization can be inferred directly from sequence, linking functional discovery, evolutionary analysis, and protein design within a shared representational framework.

## Introduction

Proteins can maintain related biological functions despite extensive divergence in sequence (Chothia & Lesk, 1986; Rost, 1999), structure (Todd et al., 2001) and conformational dynamics (Henzler-Wildman & Kern, 2007; James & Tawfik, 2003). This robustness reflects the fact that protein function is not determined solely by local residue identity or static tertiary geometry, but also by distributed physicochemical and topological relationships that act across multiple spatial scales (Bloom et al., 2006). Local interactions shape secondary structure and folding stability (Dill & MacCallum, 2012), whereas higher-order activities—including catalysis, molecular assembly, signalling and cytoskeletal remodelling—often depend on long-range coupling among domains, flexible linkers, interaction surfaces and intrinsically disordered regions (Frauenfelder et al., 1991; Halabi et al., 2009; Liu & Bahar, 2012; Motlagh et al., 2014; Wright & Dyson, 2015). How such distributed organizational features are encoded in protein sequences, and how they can be used for functional discovery or design, remains incompletely understood.

Recent advances in protein language modelling and structure prediction have transformed sequence-based protein analysis. Large-scale protein language models capture statistical constraints across evolutionary sequence space and can infer aspects of protein structure and function from sequence alone (Candido et al., 2026; Hayes et al., 2025; Lin et al., 2023; Rives et al., 2021). In parallel, deep-learning structure predictors have achieved near-atomic accuracy for many folded proteins and, more recently, biomolecular complexes (Abramson et al., 2024; Jumper et al., 2021). Structure-search methods applied to predicted proteomes have further expanded remote relationship detection at the scale of the known protein universe (Barrio-Hernandez et al., 2023; Bordin et al., 2023; van Kempen et al., 2023). Nevertheless, important regions of protein space remain difficult to interpret. Pairwise sequence similarity decays rapidly across deep evolutionary distances, predicted structures can be uncertain for flexible or disordered proteins, and many regulatory or cytoskeletal functions depend on context-dependent conformational states rather than a single rigid fold (del Alamo et al., 2022; Wayment-Steele et al., 2024). These limitations motivate approaches that can represent protein organization at an intermediate scale between amino acid sequence and three-dimensional structures.

We reasoned those hierarchical latent representations learned directly from protein sequences might capture such intermediate organizational features. In this view, protein function may be constrained by latent relationships that integrate local physicochemical compatibility with long-range structural and contextual coordination. These relationships are not necessarily visible as conserved motifs or directly alignable residues, nor are they always well represented by a single predicted structure. Instead, they may persist as distributed patterns that remain detectable across otherwise divergent sequences, folds and conformational environments.

To explore this possibility, we developed RELIC (Relational Encoding of Latent Information Contexts), a hierarchical sequence-based representation-learning framework that compresses proteins into context-dependent latent states while preserving information required for sequence reconstruction. RELIC separates local reconstruction from global contextual encoding through a compressed bottleneck, enabling sequence-derived representations to be analyzed continuously or discretized into a vocabulary of latent protein states, which we term ProtWords. The model is trained without explicit structural supervision, allowing us to test whether structural topology, functional relationships and generative constraints can emerge from sequence-derived latent organization alone.

Using RELIC, we investigate protein organization across three scales. First, we show that hierarchical bottleneck representations recover topology-associated relationships and improve remote fold-level retrieval in low-similarity regimes. Applied to the human proteome, RELIC prioritizes previously uncharacterized ciliary and microtubule-associated proteins, including C7orf57/ADMAP1, which we identify as a microtubule-associated protein required for normal sperm axonemal organization and motility in mice. Second, discretization of RELIC latent space into ProtWords reveals lineage-associated patterns of proteome organization and identifies recurrent physicochemical environments reused across divergent protein families. Third, autoregressive sampling in ProtWord space generates sequence-divergent cofilin-like proteins that retain experimentally validated F-actin disassembly and severing activity. Together, these results establish hierarchical latent representations as a framework for linking remote functional discovery, evolutionary proteome organization and protein design.

## Results

### Hierarchical bottlenecks promote topology-aware latent representations

To capture protein organization across multiple spatial scales, we developed RELIC, a hierarchical sequence representation framework that separates local sequence reconstruction from long-range contextual encoding. The architecture compresses protein sequences into a low-resolution latent bottleneck (downsampled to L/4) and reconstructs full-resolution sequences through a symmetric decoder (Fig. 1A). Skip connections between encoder and decoder preserve residue-level information locally, reducing the burden on the bottleneck to encode short-range sequence patterns. This design biases the deepest latent representations toward distributed, long-range relationships rather than local sequence similarity.

**Figure 1.**
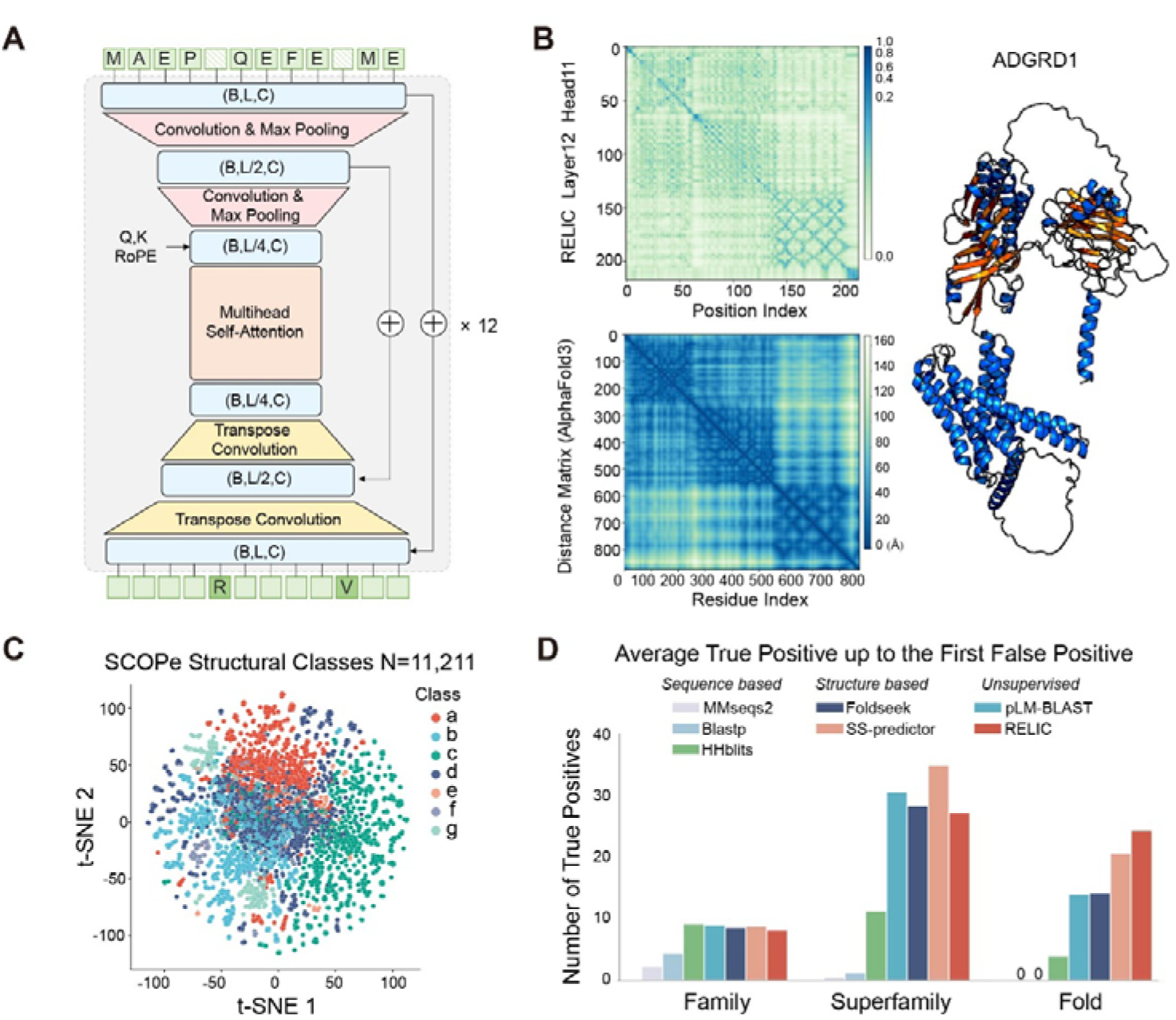
RELIC learns hierarchical protein representations associated with structural topology and improved remote fold retrieval. **(A)** Architecture of RELIC. The model employs a hierarchical convolutional encoder–decoder framework with progressive sequence downsampling through convolution and max-pooling operations to generate multiscale latent representations of protein sequences. Long-range contextual dependencies are modeled using a multi-head self-attention bottleneck operating at the coarsest latent resolution with rotary positional embeddings (RoPE) applied to query and key projections. Decoder-side transpose convolutions progressively reconstruct sequence representations, while skip connections preserve local sequence details across scales. The architecture is stacked for 12 layers and trained using a masked language modeling (MLM) objective. Tensor dimensions are denoted as batch size (B), sequence length (L), and channel dimension (C). Additional implementation details are provided in Methods. **(B)** Attention map from RELIC layer 12 head 11 compared with the corresponding AlphaFold3-predicted structural organization for ADGRD1. The upper panel shows the self-attention score matrix across latent sequential positions at the coarsest resolution of the RELIC encoder. The lower panel shows the corresponding residue–residue distance matrix derived from the AlphaFold3-predicted structure. The predicted protein structure is shown at right. Similar block-like and long-range interaction patterns are observed between the latent attention representation and the structural distance matrix. **(C)** t-SNE projection of the all-versus-all RELIC latent similarity matrix for 11,211 proteins from the SCOPe database. Points are colored according to SCOPe structural class annotations. **(D)** Remote homology retrieval on SCOPe evaluated using the average number of true positives recovered before the first false positive at the family, superfamily, and fold levels. Compared methods include sequence/profile-based approaches (MMseqs2, BLASTp, HHblits), protein language model-based retrieval (pLM-BLAST), structure-supervised representation methods (SS-predictor and Foldseek), and RELIC.

Without any explicit structural supervision, RELIC learns representations that reflect higher-order protein organization directly from primary sequences. Local structural signals, such as secondary-structure features, are still preserved (Fig. S1A), while latent attention maps exhibit block-like and long-range interaction patterns that are consistent with residue–residue distance structures observed in predicted protein models (Fig. 1B, Fig. S1B). Although attention is not a direct physical contact map, these patterns suggest that the compressed latent space retains information relevant to domain-scale topology and global structural organization.

To further investigate whether these latent representations encode meaningful structural relationships, we performed all-versus-all latent alignment across 11,211 SCOPe structural domains (Chandonia et al., 2022) using a modified Smith–Waterman algorithm (see Materials and Methods). The resulting similarity structure, visualized in embedding space, shows partial but clear organization according to major SCOPe structural classes (Fig. 1C), indicating that RELIC captures signals related to global fold architecture even when trained solely on sequence data.

We next benchmarked RELIC against established sequence-, language model-, and structure-based methods (Fig. 1D; Fig. S1C–F). RELIC outperformed conventional sequence-alignment approaches, including BLASTp (Altschul et al., 1990) and MMseqs2 (Steinegger & Söding, 2017), and profile-based approach HHblits (Remmert et al., 2012), in remote homology detection. Structure-supervised methods and approaches relying on residue-level representations, including SS-predictor (Liu et al., 2024), Foldseek (van Kempen et al., 2024) and pLM-BLAST (Kaminski et al., 2023), retained higher sensitivity at the Family and Superfamily levels, where local sequence conservation remains informative. However, this advantage diminished at deeper evolutionary distances. At the Fold level, where primary sequence similarity is largely lost, RELIC achieved the strongest retrieval performance, surpassing both Foldseek and SS-predictor especially in high-rank sensitivity. These findings suggest that hierarchical latent representations preferentially preserve distributed topological relationships conserved across deep evolutionary divergence.

### RELIC prioritizes remote microtubule-associated proteins in the human proteome

We next applied RELIC to the human proteome to determine whether latent representations could uncover functional relationships not detectable by conventional sequence or structural similarity. For each human protein query, we compared the top-ranked relationship identified by RELIC with the top-ranked relationship identified by Foldseek and evaluated the structural similarity of each retrieved pair using TM-score. Across all human queries, RELIC and Foldseek showed partially overlapping but distinct retrieval behavior, with RELIC preferentially improving matches in low-sequence-identity regimes (Fig. 2A–C). This difference was most pronounced within the sequence twilight zone, defined here as retrieved pairs with less than 30% sequence identity, where RELIC identified a larger fraction of higher-TM-score relationships than Foldseek (Fig. 2B–C). RELIC also performed favorably among proteins whose retrieved structures had lower average AlphaFold confidence, suggesting that latent representations can retain useful organizational information in regions where coordinate-based comparison is less reliable (Fig. 2D).

**Figure 2.**
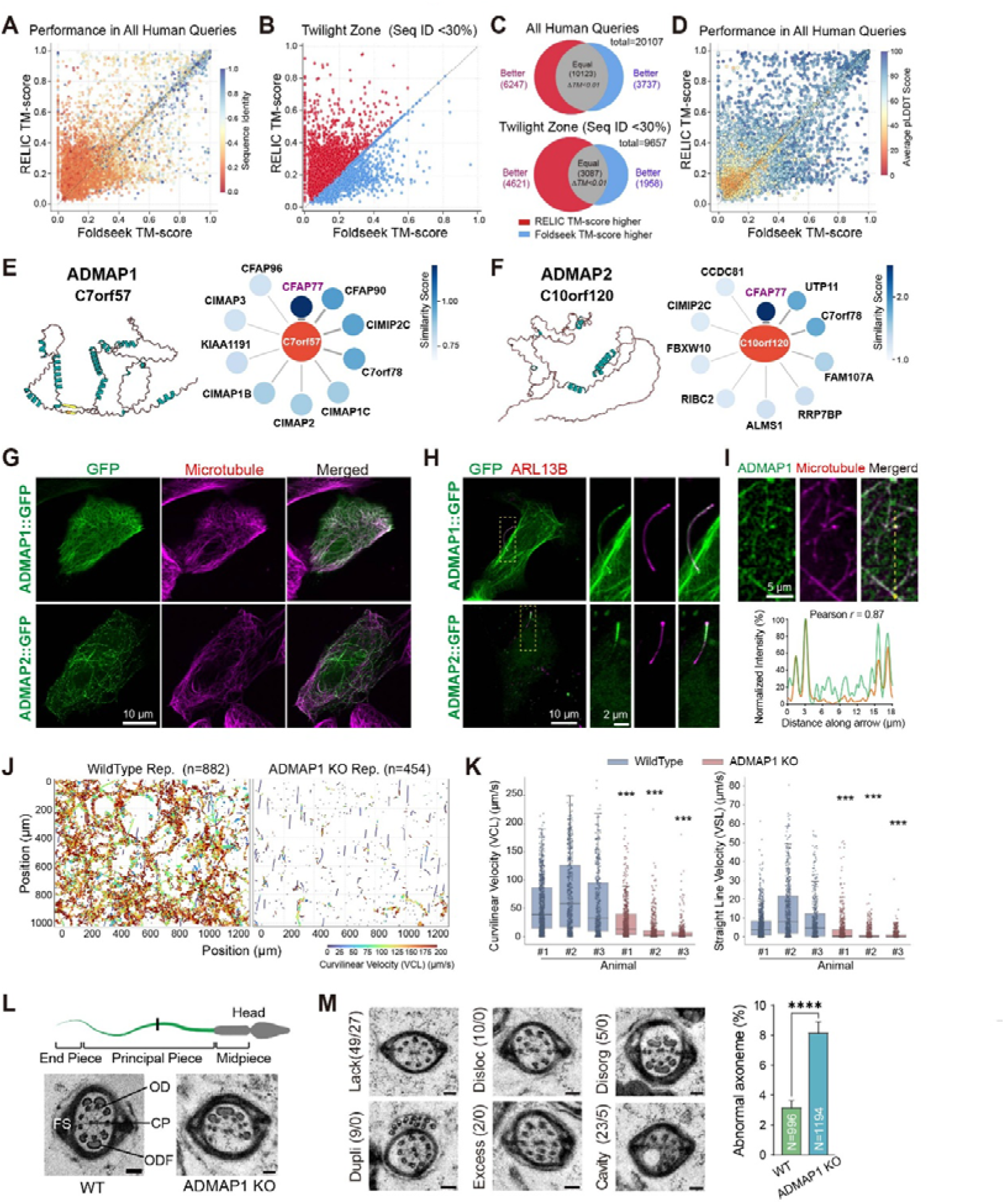
RELIC identifies remote functional relationships beyond sequence similarity and enables functional characterization of previously unannotated microtubule-associated proteins. **(A)** Pairwise comparison of structural retrieval performance between RELIC and Foldseek across all human protein queries. For each query protein, the top-ranked hit returned by each method was structurally aligned to the query using TM-score. Each point represents a query protein, with the x– and y-axes corresponding to the TM-scores of the top Foldseek and RELIC hits, respectively. Points are colored by the sequence identity between the query and the higher-scoring retrieved structure. **(B)** Structural retrieval performance within the sequence twilight zone (sequence identity <30%). Query proteins for which RELIC or Foldseek retrieved the higher-scoring structural match are highlighted. **(C)** Venn diagrams summarizing overlap between query proteins for which RELIC or Foldseek retrieved the higher-scoring structural match across all human protein queries and within the sequence twilight zone (sequence identity <30%). Structural similarity was evaluated using TM-score between each query protein and the top-ranked structure retrieved by each method. Retrievals with absolute TM-score differences smaller than 0.01 (ΔTM < 0.01) were classified as equivalent. **(D)** Structural retrieval performance across all human protein queries colored by the average AlphaFold pLDDT score of the retrieved structure from the method achieving the higher TM-score. **(E)** RELIC similarity network centered on C7orf57/ADMAP1. Predicted relationships connect C7orf57/ADMAP1 with multiple centrosomal and ciliary proteins, including CFAP77, CFAP90, CIMAP family proteins. Protein structure of C7orf57 is shown at left. **(F)** RELIC similarity network centered on C10orf120/ADMAP2. Predicted relationships include multiple centrosomal and ciliary-associated proteins, including CFAP77 and CIMIP2C. Protein structures is shown at left. **(G)** Fluorescence imaging of GFP-tagged ADMAP1 and ADMAP2 expressed in HeLa cells. Microtubules were visualized by rhodamine–taxol staining. Representative merged and single-channel images are shown. Scale bar, 10 μm. **(H)** Fluorescence imaging of GFP-tagged ADMAP1 and ADMAP2 in IMCD3 cells co-expressing mCherry-ARL13B as a primary cilium marker. Insets show enlarged views of ciliary localization. Scale bars, 10 μm and 2 μm. **(I)** Line-scan analysis of ADMAP1and microtubule fluorescence intensity along individual microtubules. Pearson correlation coefficient is indicated. Scale bar, 5 μm. **(J)** Representative sperm motility trajectories from wild-type and ADMAP1 knockout mice. Tracks are colored by curvilinear velocity (VCL). **(K)** Quantification of sperm curvilinear velocity (VCL) and straight-line velocity (VSL) in wild-type and ADMAP1 knockout animals. Each point represents an individual sperm track. Statistical significance was evaluated using two-sided Mann–Whitney U tests comparing pooled wild-type samples with each individual knockout sample, followed by Benjamini–Hochberg correction for multiple comparisons. **P < 0.001. **(L)** Schematic of sperm flagellar regions and representative transmission electron microscopy images of wild-type and ADMAP1 knockout sperm axonemes. OD, outer dynein arm; ODF, outer dense fiber; CP, central pair; FS, fibrous sheath. **(M)** Representative axonemal ultrastructural abnormalities observed in C7orf57 knockout sperm, including absence of axonemal structures, duplication defects, excess microtubule assemblies, axonemal disorganization, and cavity formation. Numbers in parentheses indicate the frequency of each defect observed in knockout and wild-type sperm (KO/WT). Quantification of abnormal axoneme frequency is shown at right. Error bars indicate standard errors (SE) estimated by binomial distribution. Statistical significance was evaluated using a chi-square test. x^2^ = 23.38, ****P < 0.0001.

Among the highest-confidence RELIC predictions, the previously uncharacterized human open reading frame C7orf57 formed a latent neighborhood enriched for ciliary and microtubule-associated proteins, including CFAP77, CFAP90 and CIMAP-family proteins (Fig. 2E). C7orf57 was not prioritized by standard sequence search and lacked a sufficiently confident structural model for robust coordinate-based comparison, making it a representative case in which latent relationships could provide functional information not readily obtained from existing annotation pipelines. Based on its RELIC-defined association with microtubule and ciliary proteins, we refer to C7orf57 here as ADMAP1, for association-defined microtubule-associated protein 1.

We experimentally tested this prediction by examining ADMAP1 localization and microtubule association. In HeLa cells, GFP-tagged ADMAP1 localized along the microtubule cytoskeleton (Fig. 2G). In ciliated IMCD3 cells, ADMAP1 was also detected along ARL13B-positive primary cilia (Fig. 2H). Microtubule depolymerization by nocodazole disrupted the filamentous localization of ADMAP1, indicating that its cellular distribution depends on polymerized microtubules (Fig. S2A). Consistent with this, recombinant ADMAP1 associated with polymerized microtubules in vitro, and line-scan analysis showed strong colocalization between ADMAP1 and microtubule fluorescence signals (Fig. 2I; Fig. S2B, C). These results support the prediction that ADMAP1 is a microtubule-associated protein.

To determine whether ADMAP1 has a physiological role in microtubule-based structures, we generated C7orf57 knockout mice using CRISPR–Cas9 genome editing (Fig. S2D). Loss of C7orf57 reduced endogenous C7orf57 signal in sperm flagella (Fig. S2E). Homozygous knockout males displayed severe defects in sperm motility, with marked reductions in curvilinear velocity and straight-line velocity across independent knockout animals (Fig. 2J, K; Fig. S2F). Transmission electron microscopy of sperm flagella revealed frequent axonemal abnormalities in knockout sperm, including loss of axonemal structures, microtubule disorganization, duplicated or excess microtubule assemblies and cavity formation (Fig. 2L, M). Quantification confirmed a significant increase in abnormal axonemal cross-sections in C7orf57 knockout sperm compared with wild-type controls (Fig. 2M). Together, these data identify ADMAP1/C7orf57 as a microtubule-associated protein required for normal sperm axonemal organization and motility.

We further examined whether other proteins in the same latent neighborhood showed related cellular behavior. C10orf120, which also clustered with ciliary and microtubule-associated proteins in RELIC space, localized to cytoplasmic microtubules and ARL13B-positive cilia when expressed in cells (Fig. 2F–H). We refer to this protein here as ADMAP2. Although additional physiological studies will be required to define its endogenous function, this independent localization result supports the ability of RELIC neighborhoods to prioritize previously uncharacterized proteins with related cellular properties. Collectively, these findings show that RELIC can nominate functional protein relationships in low-identity and structurally uncertain regions of the human proteome, enabling experimental discovery of previously unannotated proteins.

### Discrete ProtWords capture context-dependent protein organizational states

We next asked whether the continuous RELIC latent space could be represented as a discrete vocabulary of reusable protein states. To this end, we trained a vector-quantized autoencoder (VQ-VAE) (Van Den Oord & Vinyals, 2017) that maps compressed RELIC representations to a codebook of 8,192 discrete latent tokens, which we term ProtWords (Fig. 3A). Because each ProtWord is decoded in the context of surrounding latent states, we considered ProtWords not as fixed sequence motifs, but as context-dependent latent units whose amino acid realization may vary across proteins.

**Figure 3.**
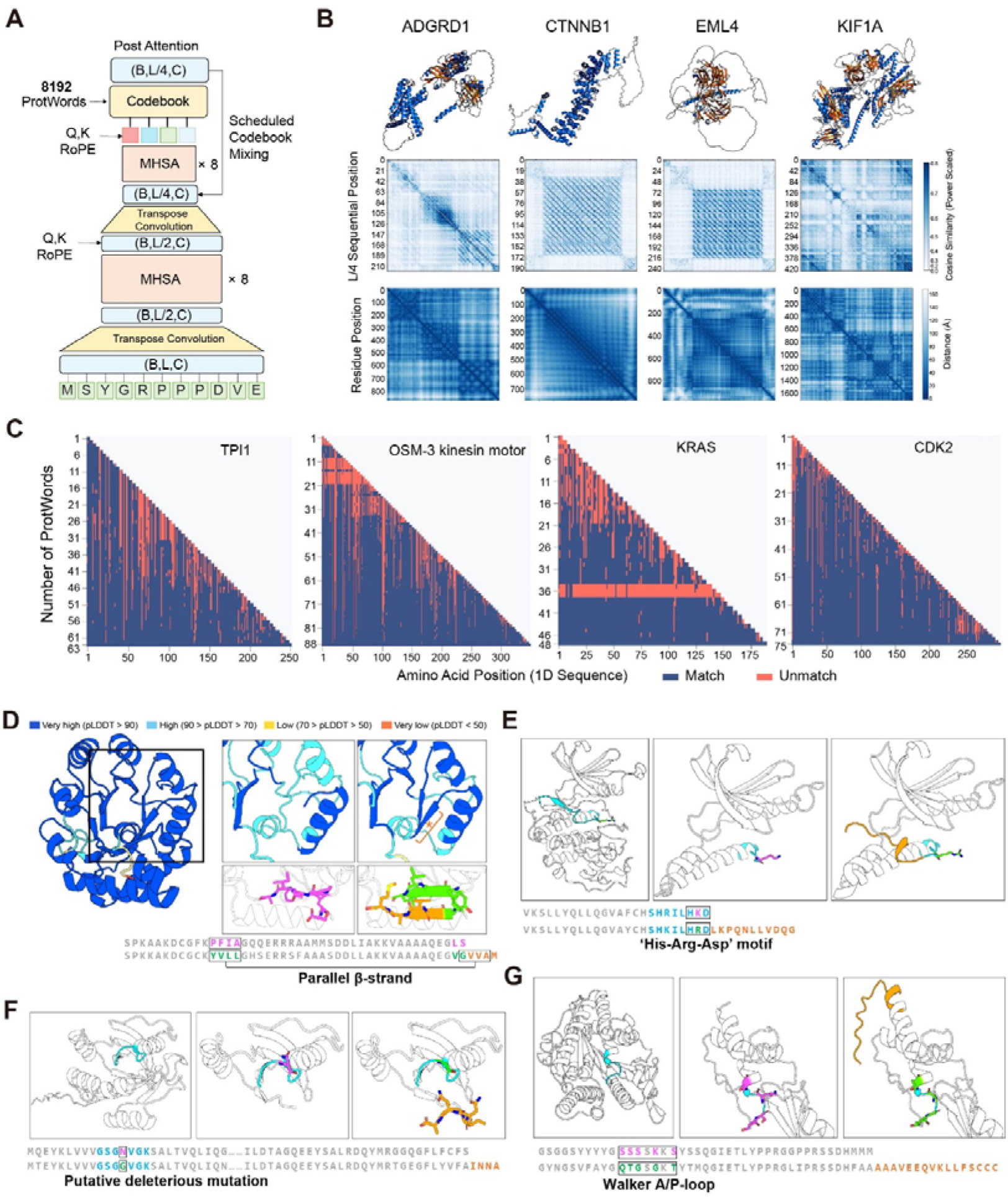
Hierarchical protein discretization reveals emergent structural organization and interpretable ProtWord representations. (**A**) Architecture of the ProtWord tokenizer. Protein sequences are progressively compressed through hierarchical convolutional and attention-based encoding layers into discrete latent representations using an 8,192-entry codebook. Rotary positional embeddings (RoPE), multi-head self-attention (MHSA), and transpose convolutions are incorporated throughout the hierarchical encoder-decoder framework. During training, scheduled codebook mixing was applied using a curriculum-learning strategy in which the sequence decoder progressively transitioned from decoding mixed continuous–discrete latent representations to decoding exclusively from discrete codebook embeddings. B, batch size; L, sequence length; C, channel dimension. (**B**) Representative examples of ProtWord organization across proteins with distinct structural architectures, including ADGRD1, CTNNB1, EML4, and KIF1A. Top panels show predicted protein structures. Middle panels show pairwise cosine similarity matrices between compressed latent representations at sequential resolution L/4. Bottom panels show corresponding AlphaFold3 residue distance matrices. (**C**) Progressive sequence realization during autoregressive ProtWord decoding for representative proteins including TPI1, the OSM-3 kinesin motor domain, KRAS, and CDK2. Each row represents the intermediate decoded sequence generated after sequential addition of ProtWords. Blue indicates amino acid positions matching the final decoded protein sequence, whereas red indicates mismatched positions. Continuous horizontal mismatch patterns arise from insertion or deletion events that shift downstream sequence alignment during intermediate decoding steps. (**D**) Example of ProtWord correspondence to local structural motifs within a parallel β-strand region. Protein structures are colored by AlphaFold pLDDT confidence scores. Insets show local structural changes associated with progressive ProtWord decoding. Bottom panels illustrate sequence realization dynamics during ProtWord addition, with colors corresponding to structural highlights above. Magenta indicates residues present in the intermediate decoded sequence, green indicates residues in the final decoded sequence, and orange indicates residues newly introduced following addition of a ProtWord. The starred structural segment in the upper inset corresponds to the newly introduced sequence region highlighted in orange below. (**E**) Example of a ProtWord associated with a catalytic loop containing the conserved His–Arg–Asp (HRD) motif. Structural context and progressive sequence realization during ProtWord decoding are shown. The catalytic loop is highlighted in cyan, the boxed residues indicate the HRD motif, residues present in the intermediate decoded sequence are shown in magenta, and residues newly introduced following ProtWord addition are shown in orange. (**F**) Example of a ProtWord associated with the P-loop region of a small GTPase. Structural context and progressive sequence realization during ProtWord decoding are shown. The Walker P-loop motif is highlighted in cyan, residues present in the intermediate decoded sequence are shown in magenta, and residues newly introduced following ProtWord addition are shown in orange, and residues in the final decoded sequence are shown in green. (**G**) Example of a ProtWord associated with a Walker A/P-loop motif. Structural context and progressive sequence realization during ProtWord decoding are shown. The Walker A/P-loop motif is highlighted in cyan. Residues present in the intermediate decoded sequence are shown in magenta, residues newly introduced following ProtWord addition are shown in orange, and residues in the final decoded sequence are shown in green. Sequence segments corresponding to intermediate and final decoded states are shown below.

We first examined whether these discrete representations preserve information about global protein organization. Conceptually, we anticipated that ProtWords could serve as descriptors of higher-order spatial environments. Based on this rationale, spatially proximal regions within a folded protein should share similar contextual features, allowing pairwise similarities between these latent states to reflect the protein’s overall structural architecture. To evaluate this hypothesis, we decoded ProtWord sequences for each protein into intermediate latent representations before full amino acid reconstruction and computed pairwise similarity between latent positions. Across proteins with distinct architectures, including ADGRD1, CTNNB1, EML4 and KIF1A, ProtWord-derived similarity matrices recovered block-like organization and long-range relationships that paralleled domain-scale features in AlphaFold3-predicted distance maps (Fig. 3B, S4D). These results indicate that compact discrete latent representations retain information associated with structural topology as a whole seqeunce, despite operating at a reduced sequence resolution.

To determine whether amino acid reconstruction from ProtWords depends on local token identity alone or on broader latent context, we progressively extended the ProtWord input context and monitored how the decoded amino acid sequence changed after each added token. Across representative proteins, including TPI1, an OSM-3 motor domain, KRAS and CDK2, newly added downstream ProtWords altered not only adjacent residues but also previously decoded upstream regions (Fig. 3C). This behavior indicates that sequence realization is globally conditioned rather than strictly left-to-right or locally specified. In several examples, context extension corrected register shifts, stabilized secondary-structure pairing and improved local packing. For example, addition of downstream ProtWords promoted compatible sequence realization across a parallel β-strand region, with corresponding increases in predicted local confidence in the reconstructed structure (Fig. 3D).

Context-dependent reconstruction also affected conserved functional regions. In kinase-like proteins, expansion of the ProtWord context restored residues corresponding to the His–Arg–Asp catalytic-loop motif (Fig. 3E). In small GTPase and kinesin-family examples, extended context improved reconstruction of P-loop or Walker A–like nucleotide-binding motifs (Fig. 3F, G). These examples suggest that ProtWords can encode reusable latent states associated with structural and functional environments, while the final amino acid sequence is determined by broader context.

Thus, discretization of RELIC latent space yields ProtWords, a compact vocabulary of context-dependent protein states that captures structural topology, local packing and functional-site organization. Rather than representing fixed sequence motifs, ProtWords encode distributed structural constraints that guide amino acid realization across both local and long-range protein environments.

### Proteome-wide ProtWord usage captures lineage-associated protein organization

Building upon these insights into individual proteins, we expanded our analysis to compare protein organization at the scale of entire proteomes. We encoded proteins from 54 representative species into the shared ProtWord vocabulary and quantified proteome-wide token usage patterns across archaea, bacteria, protists, fungi, plants and metazoans (Fig. 4A; Fig. S3). Within each species, ProtWord usage was heterogeneous across the proteome, indicating that the vocabulary captures diverse protein organizational states rather than species identity alone.

**Figure 4.**
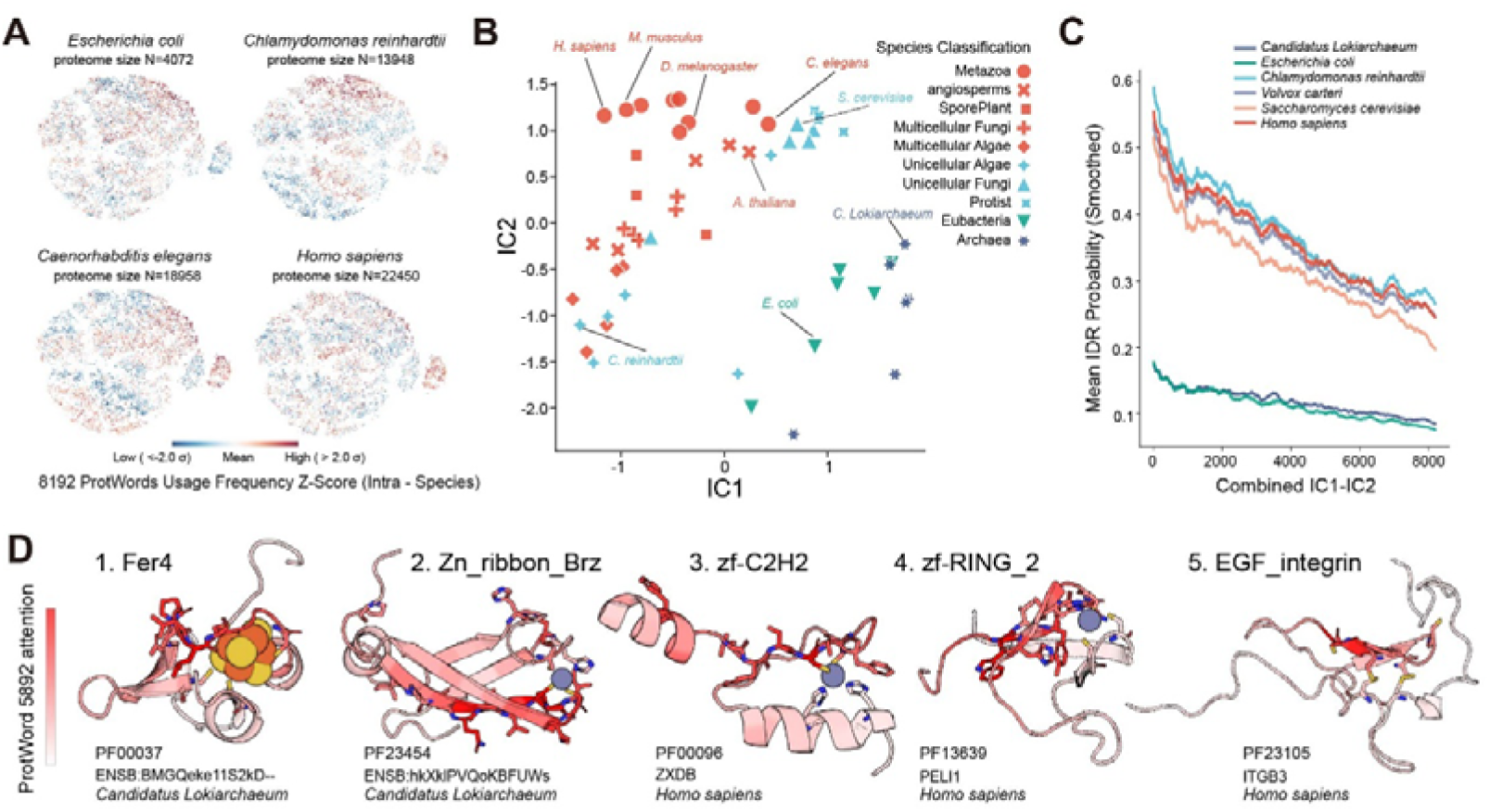
ProtWord usage patterns capture species-level organizational signatures and recurrent structural motifs. (**A**) UMAP projections of proteome-wide ProtWord usage profiles from representative species, including *Escherichia coli*, *Chlamydomonas reinhardtii*, *Caenorhabditis elegans*, and *Homo sapiens*. Points represent proteins colored by intra-species ProtWord usage frequency Z-scores. Proteome sizes are indicated for each species. (**B**) ICA projection of species-level normalized ProtWord usage profiles across representative organisms spanning archaea, bacteria, protists, fungi, plants, and metazoans. Species are colored and shaped according to taxonomic classification. (**C**) Mean intrinsic disorder region (IDR) propensity across the ProtWord ICA landscape for representative species. ProtWords were ordered along the combined IC1–IC2 trajectory (IC1-IC2), revealing systematic differences in disorder-associated ProtWord usage across taxa. (**D**) Representative physicochemical microenvironments associated with high-attribution ProtWords. Examples include iron-sulfur cluster coordination environments, zinc ribbon folds, C2H2 zinc finger domains, RING finger domains, and EGF-integrin-associated architectures. Residue-level integrated gradients attribution scores for ProtWord 5892 are mapped onto protein structures and highlighted in red. Representative proteins and corresponding species are indicated below each structure.

We then summarized species-level ProtWord usage profiles using independent component analysis. This analysis separated prokaryotic and eukaryotic proteomes and further distributed multicellular eukaryotes, unicellular eukaryotes, fungi, plants and metazoans along a continuous latent trajectory (Fig. 4B; Fig. S5A, B). These patterns indicate that the relative usage of discrete latent protein states differs systematically across major evolutionary groups.

We next examined whether this species-level organization was associated with biophysical properties of proteins. Ordering ProtWords along the combined IC1–IC2 axis revealed a strong association with predicted intrinsic disorder. ProtWords enriched in eukaryotic proteomes were associated with higher local disorder propensity, whereas ProtWords enriched in bacterial and archaeal proteomes were associated with lower predicted disorder (Fig. 4C; Fig. S5C,D). Gene Ontology enrichment analysis further linked eukaryote-associated ProtWords to regulatory, developmental and signaling-related annotations, whereas prokaryote-associated ProtWords were enriched for catalytic, transport and metabolic functions (Fig. S4A–C). These results suggest that ProtWord usage captures a broad evolutionary shift from compact enzymatic and metabolic protein environments toward expanded disordered and regulatory protein contexts in eukaryotic proteomes.

To test whether individual ProtWords correspond to recurrent physicochemical environments across divergent proteins, we examined ProtWord 5892, one of the strongly weighted ProtWords along the eukaryote-associated region of the ICA landscape (Ranking No. 347). Proteins containing this ProtWord spanned diverse species, sequences and global folds, yet residue-level attribution consistently highlighted cysteine-rich coordination environments. Representative examples included iron–sulfur cluster-binding ferredoxin-like proteins, zinc-ribbon folds, C2H2 zinc fingers, RING-finger domains and EGF–integrin-associated disulfide-rich architectures (Fig. 4D). Therefore, individual ProtWords can mark reusable local physicochemical environments that appear in distinct structural and evolutionary contexts. Together, these analyses show that proteome-wide ProtWord distributions capture lineage-associated organizational signatures and that specific ProtWords can represent recurrent biochemical environments across protein space.

### Latent-space generation preserves foldable protein organization

Having established that ProtWords capture fundamental organizational principles, we explored whether this discrete vocabulary could support generative protein design. Existing protein design strategies commonly operate either by first specifying a backbone (Watson et al., 2023) and then optimizing sequence (Dauparas et al., 2022), or by directly sampling amino acid sequences from sequence-based language models (Ferruz et al., 2022; Madani et al., 2023; Nijkamp et al., 2023). In contrast, RELIC enables generation at an intermediate latent scale: an autoregressive model samples ProtWord trajectories, and the hierarchical decoder then realizes these trajectories as amino acid sequences in a context-dependent manner (Fig. 5A). This design allows global organizational states and local sequence features to be coupled during generation rather than optimized as separate steps.

**Figure 5.**
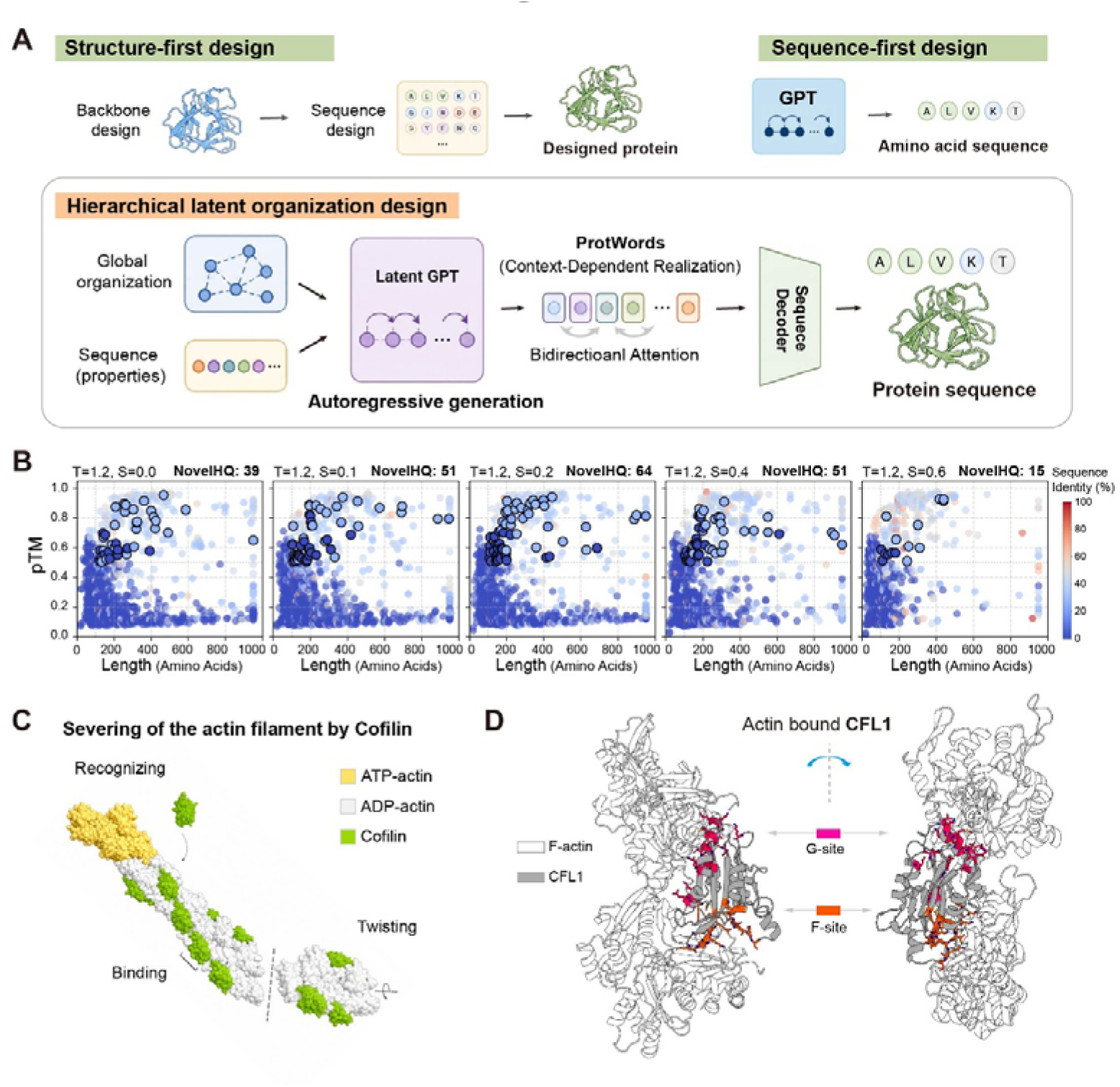
Hierarchical latent organization design enables generation of structurally diverse actin-remodeling proteins. (**A**) Comparison of representative protein design paradigms. Structure-first approaches generate protein sequences from predefined structural scaffolds, whereas sequence-first approaches directly model amino acid sequences autoregressively. RELIC hierarchical latent organization design instead operates in a latent semantic space, integrating higher-order organizational features and sequence-level properties through autoregressive ProtWord generation followed by sequence decoding. (**B**) Structural quality assessment of generated proteins across different sampling temperatures (T) and latent similarity constraints (S). Predicted TM-score (pTM) is plotted against protein length for generated sequences. During autoregressive generation, candidate ProtWords were restricted according to cosine similarity to the highest-probability ProtWord in latent space. Points are colored by sequence identity to the nearest UniRef100 neighbor. NovelHQ indicates the number of generated proteins with high predicted structural confidence and low sequence similarity to known proteins. (**C**) Schematic model of cofilin-mediated actin filament severing. Cofilin preferentially recognizes aged ADP-actin filament conformations, followed by cooperative filament binding, filament twisting, and eventual filament severing. ATP-actin, ADP-actin, and cofilin molecules are indicated by color. (**D**) AlphaFold3 predicted structural model of the CFL1–actin binding interface. The ribbon representation depicts natural human cofilin (CFL1, gray) complexed with the F-actin filament (white). Crucial functional interfaces are color-coded to illustrate the distributed nature of the interaction: the G-actin binding site (G-site) is highlighted in magenta, and the F-actin binding site (F-site) is highlighted in orange. This complex architecture establishes the structural baseline for target-site remodeling and design.

To implement this strategy, we trained an autoregressive transformer, termed Latent GPT, to model transitions between ProtWords. Generated ProtWord sequences were decoded into amino acid sequences using the VQVAE decoder and evaluated by ESMFold. Because sequence realization depends on distributed latent context, generation within ProtWord space constrains both global organization and local sequence environments simultaneously.

Structural analysis demonstrated that de novo generated proteins retained substantial predicted folding confidence and core packing integrity (Fig. 5B; Fig. S6A–C). In contrast, randomization of either ProtWord order or decoded amino acid sequences disrupted predicted structural organization and sharply reduced folding confidence (Fig. S6D). These findings indicate that autoregressive latent-space sampling preserves distributed organizational constraints required for coherent protein folding.

### Latent-space generation produces functional actin-remodeling proteins

Moving beyond computational predictions, we sought to determine whether ProtWord-based latent generation could yield proteins with experimentally measurable biochemical activity. We focused on the actin-depolymerizing factor/cofilin family, whose filament-severing activity requires coordinated interactions across multiple actin-binding surfaces and conformational remodeling of F-actin (Fig. 5C, D; Fig. S7A) (Galkin et al., 2011; Huehn et al., 2020). Cofilin engages actin through a sequential induced-fit mechanism involving substantial conformational rearrangements (Tanaka et al., 2018), while its activity is further regulated by phosphorylation, N-terminal allosteric cues, local pH, and partner proteins including coronin and AIP1 (Brieher et al., 2006; Oosterheert et al., 2025; Pope et al., 2004; Sexton et al., 2024). Unlike functions that can be approximated by a single catalytic residue or static binding site, cofilin activity depends on distributed features of the ADF-H fold, including the G-actin-binding surface (G-site), F-actin-binding interface (F-site), N-terminal regulatory region and hydrophobic core (Fig. 5D). This made cofilin a stringent test of whether latent-space generation can preserve functionally relevant organizational constraints while allowing substantial sequence diversification.

We fine-tuned Latent GPT on ADF/cofilin-family ProtWord representations and decoded generated latent trajectories into amino acid sequences, which we refer to as pwCofilins. From 100 generated sequences, 57 candidates were selected on the basis of sequence divergence from native cofilins and predicted foldability. Most selected candidates retained predicted ADF-H-like structures, with 55 of 57 showing ESMFold pTM scores greater than 0.7 (Fig. S7B). We then experimentally evaluated 11 randomly selected pwCofilins in mammalian cells. All tested designs were detectably expressed, and most showed diffuse cytosolic localization without obvious insoluble aggregation or condensate formation (Fig. S7C).

Expression of the non-phosphorylatable form (S3A) of three generated proteins, pwCofilin7, pwCofilin14 and pwCofilin90, induced pronounced remodeling of the actin cytoskeleton in cells (Fig. 6A). Quantification of phalloidin-stained cells showed that these designs increased the fraction of cells with severed or disrupted F-actin structures relative to wild-type CFL1, with pwCofilin7 and pwCofilin14 reaching activity levels comparable to constitutively active CFL1(S3A) in this assay (Fig. 6B). Thus, a subset of generated cofilin-like proteins retained cellular actin-remodeling activity despite substantial sequence divergence from natural cofilins.

**Figure 6.**
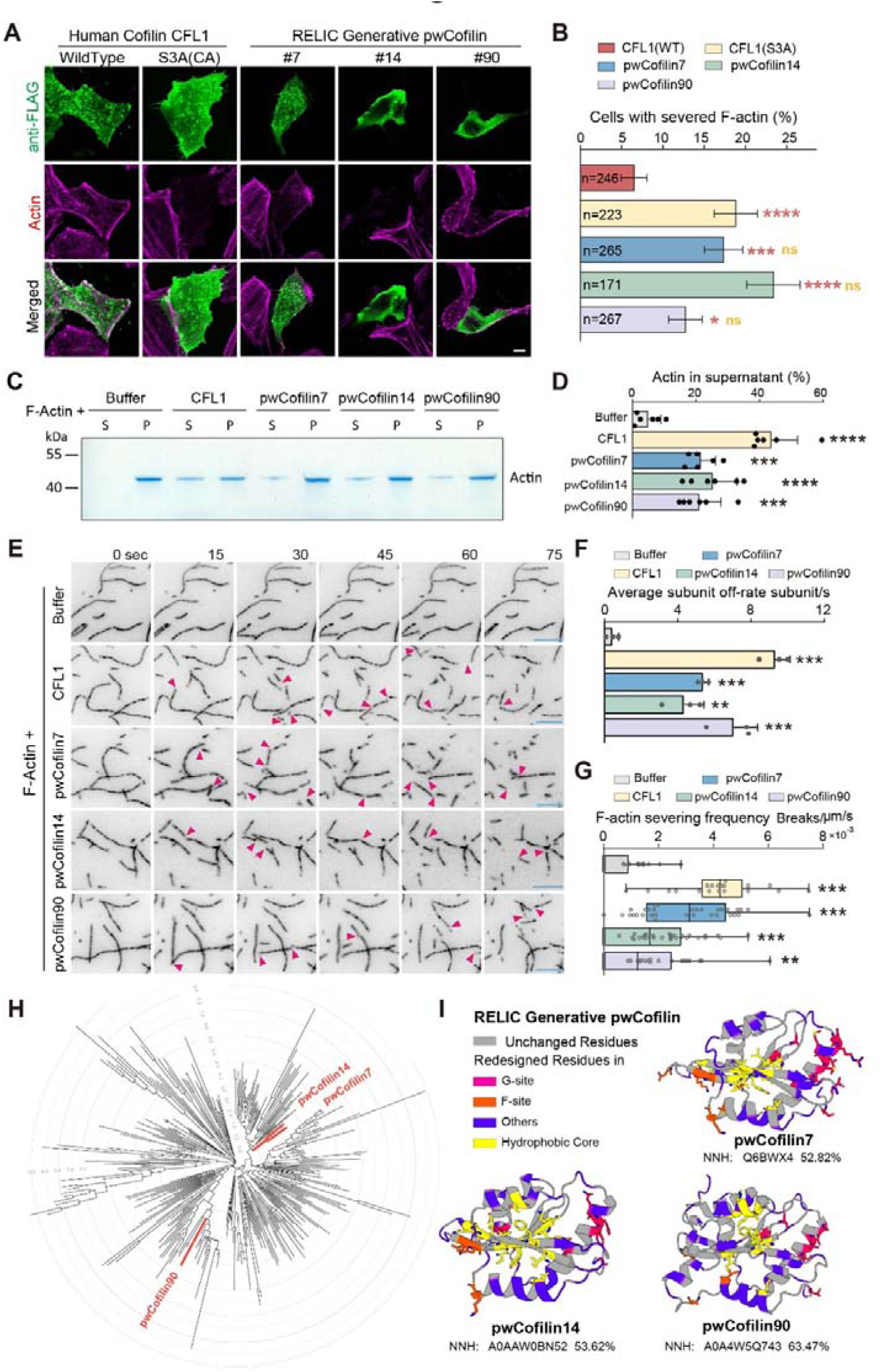
Generated cofilin-like proteins retain actin-remodeling activity despite extensive sequence and organizational divergence. (**A**) Cellular expression of FLAG-tagged human CFL1(WT), constitutively active CFL1(S3A), and generated cofilin-like proteins (pwCofilin7, pwCofilin14, and pwCofilin90) in HeLa cells. Cells were stained with anti-FLAG antibody (green) and phalloidin-labeled F-actin (magenta). Representative images are shown. Scale bar, 10 μm. (**B**) Quantification of cells exhibiting severed or disrupted F-actin structures following expression of wild-type, mutant, or generated cofilin-like proteins. Each cell was classified based on the presence or absence of severed F-actin morphology. Numbers of analyzed cells are indicated within bars. Error bars represent standard deviation estimated from a binomial distribution. Statistical significance was evaluated using chi-square tests with Benjamini–Hochberg correction for multiple comparisons. Red markers indicate statistical comparisons with CFL1(WT), while yellow markers indicate comparisons with CFL1(S3A). ns, not significant; *P < 0.05; ***P < 0.001; ****P < 0.0001. (**C**) F-actin severing assay performed with purified actin and indicated cofilin variants. Following incubation with polymerized F-actin, samples were separated into supernatant (S) and pellet (P) fractions by ultracentrifugation and analyzed by SDS–PAGE followed by Coomassie staining. Redistribution of actin from pellet to supernatant fractions reflects cofilin-mediated disruption of filamentous actin. (**D**) Quantification of actin recovered in the supernatant fraction from the F-actin severing assays shown in (C). Each point represents an independent experimental replicate. Error bars indicate standard deviation. Statistical significance was evaluated using two-sided Mann–Whitney U tests with Benjamini–Hochberg correction for multiple comparisons. **P < 0.01; ***P < 0.001; ****P < 0.0001. (**E**) Time-lapse imaging of individual actin filaments following incubation with purified CFL1(S3A), pwCofilin7, pwCofilin14, or pwCofilin90. Magenta arrowheads indicate filament breakage sites. Time is shown in seconds. Scale bar, 5 μm. (**F**) Quantification of average actin filament subunit dissociation rates following treatment with indicated cofilin variants. Data were acquired from three independent fields of view for each condition. Error bars indicate standard deviation. Statistical significance was evaluated using two-sided Mann–Whitney U tests with Benjamini–Hochberg correction for multiple comparisons relative to buffer control. **P < 0.01; ***P < 0.001. (**G**) Quantification of F-actin severing frequency normalized by filament length and time from experiments shown in (E). Data were acquired from three independent fields of view for each condition. Error bars indicate standard deviation. Statistical significance was evaluated using two-sided Mann–Whitney U tests with Benjamini–Hochberg correction for multiple comparisons relative to buffer control. ***P < 0.001. (**H**) Phylogenetic distribution of experimentally validated pwCofilins relative to natural ADF/cofilin family proteins from UniRef100. Natural cofilin-family proteins were clustered at 50% sequence identity and 80% coverage prior to tree construction to reduce redundancy. Functional generated variants (highlighted in red) formed multiple distinct peripheral branches across sequence space rather than collapsing into a single localized clade adjacent to known natural homologs, despite retaining robust actin-remodeling activity. Scale bar indicates expected amino-acid substitutions per aligned site estimated under the Q.PFAM+R6 evolutionary model. (**I**) Structural mapping of redesigned residues in representative generated cofilin-like proteins (pwCofilin7, pwCofilin14, and pwCofilin90) modeled in complex with actin using AlphaFold3. Residues differing from the nearest natural homolog (NNH) in UniRef100 are highlighted and categorized according to canonical cofilin functional regions, including the G-site (magenta), F-site (orange), hydrophobic core (yellow), and other positions (purple). Gray residues indicate positions conserved relative to the nearest natural homolog. Sequence identity to the nearest UniRef100 neighbor is indicated below each structure.

To test whether these cellular phenotypes reflected intrinsic biochemical activity, we purified recombinant CFL1, pwCofilin7, pwCofilin14 and pwCofilin90 (Fig. S7D) and examined their effects on polymerized actin in vitro. In F-actin sedimentation assays, all three pwCofilins increased the fraction of actin recovered in the supernatant relative to buffer control, consistent with filament destabilization or disassembly (Fig. 6C, D). We then directly measured filament remodeling dynamics by time-lapse total internal reflection fluorescence (TIRF) microscopy. Purified pwCofilin7, pwCofilin14 and pwCofilin90 induced visible breakage of individual immobilized actin filaments (Fig. 6E). Quantification showed increased actin subunit loss and elevated severing frequency relative to buffer control under the assay conditions tested (Fig. 6F, G). These results demonstrate that the active pwCofilins possess intrinsic F-actin disassembly and severing activity.

We next evaluated the relationship between our functional designs and the natural cofilin sequence space. Rather than clustering near a single natural ancestor or collapsing into a specific evolutionary branch, the three active pwCofilins occupied distinct peripheral lineages across the training set hierarchy (Fig. 6H). Notably, with sequence identities to their nearest natural homologs ranging from only 52.8% to 63.5% (Fig. 6I), these designs represent a substantial departure from simple local interpolation, confirming that the generative model can discover novel and diverse sequence solutions far beyond immediate training training examples. Multiple sequence alignment and structural modeling suggested that the active designs retained the overall ADF-H fold while redistributing substitutions across canonical cofilin functional regions, including the G-site, F-site and hydrophobic core (Fig. 6I; Fig. S7E). These changes indicate that actin-remodeling activity was preserved despite substantial remodeling of residues that are conserved in natural cofilins.

Together, these results show that family-conditioned ProtWord generation can produce sequence-divergent, structurally plausible cofilin-like proteins with experimentally validated F-actin remodeling activity. More broadly, they are consistent with the idea that latent-space generation can preserve distributed organizational constraints underlying a dynamic biochemical function while allowing multiple distinct sequence solutions.

## Discussion

This study demonstrates that higher-order protein organization and functional manifolds can emerge intrinsically from primary sequences through hierarchical latent representations. While traditional sequence alignment and coordinate-based comparisons remain foundational (Altschul et al., 1990; Steinegger & Söding, 2017; van Kempen et al., 2023), their sensitivity progressively declines within the sequence twilight zone or when encountering conformational flexibility. Advanced multimodal deep learning frameworks have recently sought to transcend these boundaries by explicitly projecting sequences, 3D structures, and natural language descriptions into unified representation spaces (Liu et al., 2025; Su et al., 2025). Although this cross-modal integration offers a powerful paradigm for data synthesis, it leaves a fundamental conceptual question open: can such higher-order relationships emerge natively from sequence data alone, without explicit structural or textual curation? Our findings indicate that compressing sequence information through a hierarchical structural bottleneck preferentially isolates distributed topological relationships. This approach not only renders representations resilient against deep evolutionary divergence and conformational plasticity, but also establishes a unified framework that successfully bridges remote homology detection, functional prioritization, and generative design.

A major conceptual implication of this work is that latent protein neighborhoods can uncover hidden functional relationships where conventional annotation pipelines fail. The prioritization and subsequent experimental validation of uncharacterized microtubule-associated proteins (ADMAP1 and ADMAP2) illustrate this capability. Crucially, these functional associations eluded not only standard alignment tools and confidence-filtered structural matches, but also state-of-the-art, residue-resolution protein language models scaling up to 6B-parameters (Candido et al., 2026) (see Supplementary Materials). This divergence in performance suggests a fundamental architectural distinction: while massive, continuous pLMs excel at capturing fine-grained local semantics, their unconstrained attention mechanisms can become saturated with noise in the sequence “twilight zone.” In contrast, RELIC’s hierarchical bottleneck filters out volatile local variations, isolating the lower-resolution topological signals that dictate cellular localization and macro-molecular indexing.

Discretizing this continuous space into a ProtWord vocabulary provides a scalable framework for studying protein evolution as trajectories through discrete organizational states. Our analysis indicates that ProtWords do not function as static, deterministic sequence motifs or structural fragments; rather, they operate as context-dependent latent units. The observation that downstream token context actively remodels upstream sequence realization challenges the conventional view of linear or strictly local sequence-to-structure translation. At the proteome scale, the systematic shift in ProtWord usage across lineages mirrors a fundamental macro-evolutionary transition: moving from the compact, structured enzymatic environments of prokaryotes toward the expanded, fluid, and regulatory disordered regions characteristic of eukaryotic complexity. Furthermore, the recurrence of identical ProtWords across entirely unrelated global folds underscores a modular evolutionary principle, wherein nature achieves structural diversification through the combinatorial reuse of invariant physicochemical environments.

The generative experiments transition this framework from passive representation to active design, establishing that autoregressive sampling within a discrete latent space can sustain complex biological functions. By focusing on the ADF/cofilin family—a system governed by cooperative, distributed interaction surfaces rather than a centralized catalytic triad—we challenged the model to capture dynamic, fold-wide constraints. The successful generation of functional, sequence-divergent variants offers key methodological insights. First, it demonstrates that latent trajectories encode the necessary global syntax to ensure coherent protein folding without explicit geometric supervision. Second, the wide phylogenetic dispersion of active designs across multiple peripheral lineages proves that the generative model does not rely on local interpolation or memorize training examples. Instead, it maps and samples from the underlying functional manifold, decoupling biological activity from rigid sequence identity and permitting expansive excursions through sequence space.

Several limitations outline directions for future research. Although RELIC enhances remote relationship detection, the interpretation of specific latent states remains probabilistic and context-dependent, necessitating more systematic functional mapping across larger datasets. Additionally, RELIC lacks explicit atomic-resolution modeling of side-chain geometry, ligand states, and post-translational modifications, making it a complementary counterpart to—rather than a replacement for—coordinate-based biophysical simulations. Finally, while the generative framework proved highly effective in a family-conditioned setting, its capacity for unconstrained, fully de novo design of novel topologies requires broader experimental validation. Interfacing hierarchical latent spaces with atomic modeling and high-throughput screening will be essential to translate these organizational principles into precise molecular engineering pipelines.

In conclusion, our findings support a unified model where remote homology, proteome evolution, and functional design are not separate computational challenges, but interconnected manifestations of shared organizational constraints embedded in sequence space. Hierarchical latent spaces offer a mathematically rigorous yet biologically intuitive framework to navigate this space, viewing proteins not merely as static arrays of coordinates or string of letters, but as functional trajectories traversing an underlying evolutionary manifold.

## Biosecurity and Dual-Use Statement

RELIC is conceived as a fundamental computational framework for advancing the understanding of protein structure, evolution, and fundamental cell biology. Although its hybrid architecture and discrete vocabulary enable powerful generative capabilities for *in silico* protein design, the framework is intended exclusively for basic, translational, and therapeutic research purposes. The pre-training dataset (UniRef50) inherently encompasses a vast breadth of natural evolutionary diversity; however, RELIC has not been fine-tuned, nor does it possess the specialized zero-shot physical modeling constraints required, to intentionally design functional viral evasion mechanisms, highly potent toxins, or novel pathogens.

We acutely recognize the dual-use nature of generative protein language models. Therefore, to mitigate potential misuse, the pre-trained weights, VQ-VAE codebooks, and inference pipelines will be publicly released under **OpenRAIL++-M License**. This licensing framework explicitly prohibits the utilization of our models, derived latent vocabularies, or downstream code for the development of biological weapons, the enhancement of pathogen virulence, or any applications that pose a deliberate threat to public health and global biosecurity.

## Data and Code Availability

The source code for the RELIC framework, including the hierarchical encoder, discrete codebook, generative latent GPT, and inference scripts, is publicly available on GitHub at https://github.com/young55775/RELIC/.

Due to file size limits, all pre-trained model checkpoints and the processed datasets required to reproduce the analyses and figures in this study have been deposited on Zenodo and are freely accessible at https://doi.org/10.5281/zenodo.20461149.

## Acknowledgements

This work was supported by the National Natural Science Foundation of China (NSFC) Grants 325B2026(Z.G.), 92254306 (G.O.), 32430026 (G.O.), 32021002 (G.O.), 32270721 (Y.C.), 32400610 (Z.W.), 32270773 (W.L.), the National Key R&D Program of China Grants 2022YFA1302700 (W.L.), 2024YFA1307301 (W.L.), and TsienTang Life Science Development Fund at Tsinghua University. We are deeply grateful to for their insightful discussions and constructive feedback. We thank the Facility of Cell Imaging in the Tsinghua University Technology Center for Protein Research (especially Yuke Feng), and the Tsinghua University Laboratory Animal Resources Center (THU-LARC) (especially Yankun Yang), for their technical assistance. We also thank the Tsinghua University Cryo-EM Facility of China National Center for Protein Sciences (Beijing) for HPF and EM data collection.

## Author Contributions

Z.G. and G.O. conceived and designed this study. G.O. supervised the project. Z.W. amd S.W performed the cell biology and in vivo mouse experiments. K.X. performed the in vivo mouse experiments and analyzed the data. M.L. conducted the transmission electron microscopy (TEM) analysis. G.O. and Z.G. wrote the manuscript with contributions from all other authors.

**Figure S1.**
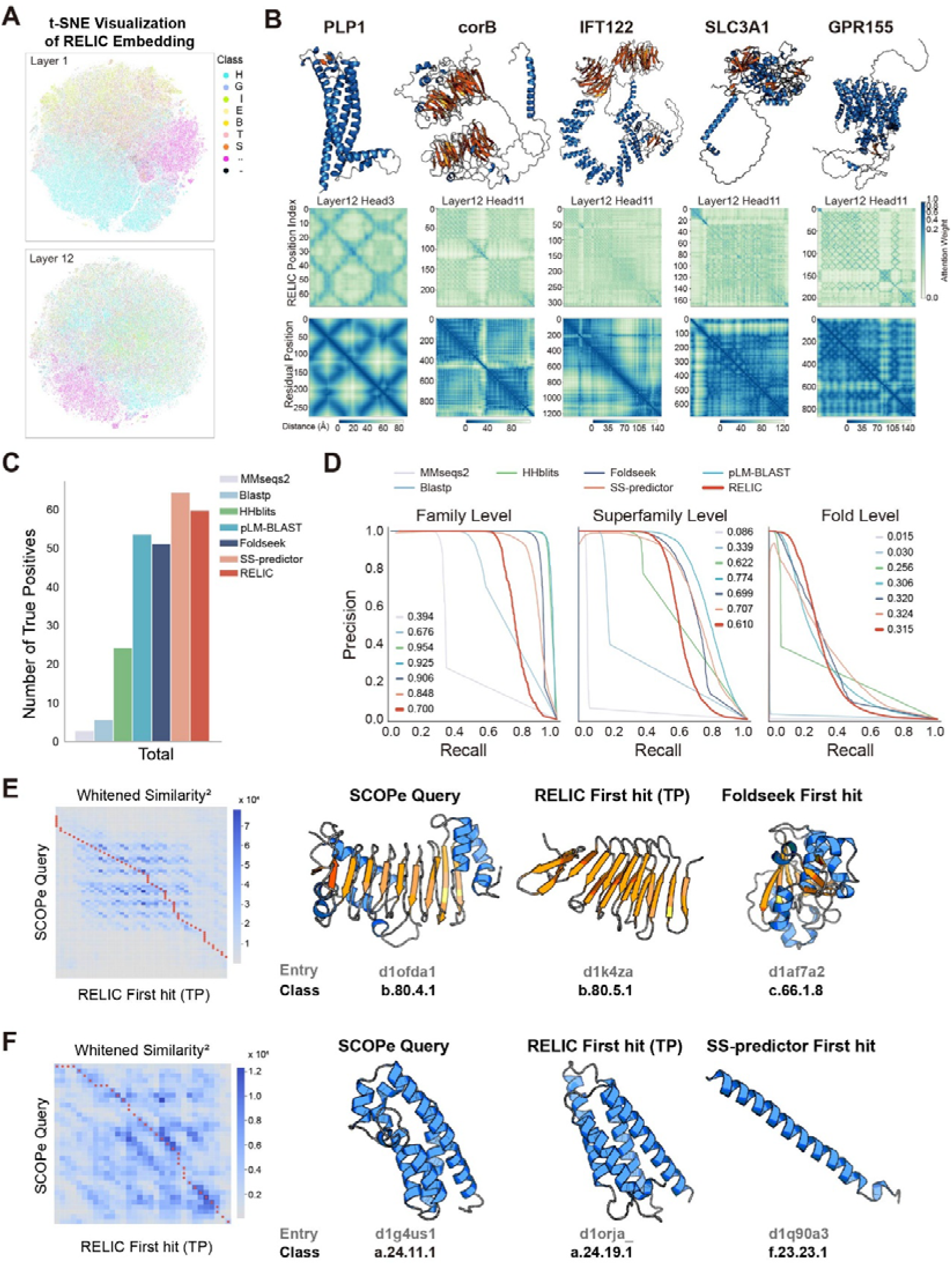
Hierarchical structural representations and benchmarking of RELIC embeddings. **(A)** t-SNE visualization of bottleneck-layer residue representations from RELIC layers 1 and 12. Embeddings were extracted from Swiss-Prot proteins with corresponding AlphaFold Database structural models and colored according to secondary structure assignments predicted using the MDAnalysis package. Early-layer representations exhibit stronger organization by local secondary structure states, whereas deeper bottleneck representations show increased mixing of higher-order structural context. **(B)** Structural interpretability of RELIC attention maps. Top row: 3D ribbon representations of five representative proteins spanning diverse topological folds (PLP1, corB, IFT122, SLC3A1, and GPR155). Middle row: Self-attention weight matrices extracted from specific heads of the deep latent layer (Layer 12, Head 3 or Head 11), with color density indicating relative attention weights. Bottom row: True ground-truth residue–residue physical distance matrices (measured in Angstroms, Å) calculated from the corresponding 3D structural coordinates. **(C)** Benchmark comparison of structural retrieval performance across SCOPe family, superfamily, and fold-level relationships using true positives identified before the first false positive (TP up to 1st FP). Methods compared include MMseqs2, BLASTp, HHblits, pLM-BLAST, Foldseek, SS-predictor, and RELIC. **(D)** Example comparison between RELIC and Foldseek retrieval trajectories for a SCOPe query protein. Left, covariance-whitened similarity matrix computed from final bottleneck-layer representations, with the optimal alignment trajectory indicated in red. Right, structural comparison between the SCOPe query, the RELIC-retrieved first hit, and the Foldseek-retrieved first hit. SCOPe entries and classifications are indicated below. **(E)** Precision-Recall curves for remote homology detection across structural hierarchies. Performance comparison of RELIC against sequence-based (MMseqs2, BLASTp, HHblits), language model-based (pLM-BLAST), and structure-based/supervised methods (Foldseek, SS-predictor) across three evolutionary depths defined by the SCOPe database: Family level (left), Superfamily level (middle), and Fold level (right). Numerical values in the legend indicate the Area Under the Precision-Recall curve (AUPRC) for each corresponding method. RELIC preferentially maintains high precision at deeper evolutionary distances (Fold level), particularly in the low-recall/high-rank sensitivity regime. **(F)** Example comparison between RELIC and SS-predictor retrieval trajectories for a SCOPe query protein. Left, covariance-whitened similarity matrix computed from final bottleneck-layer representations, with the optimal alignment trajectory indicated in red. Right, structural comparison between the SCOPe query, the RELIC-retrieved first hit, and the SS-predictor-retrieved first hit. SCOPe entries and classifications are indicated below.

**Figure S2.**
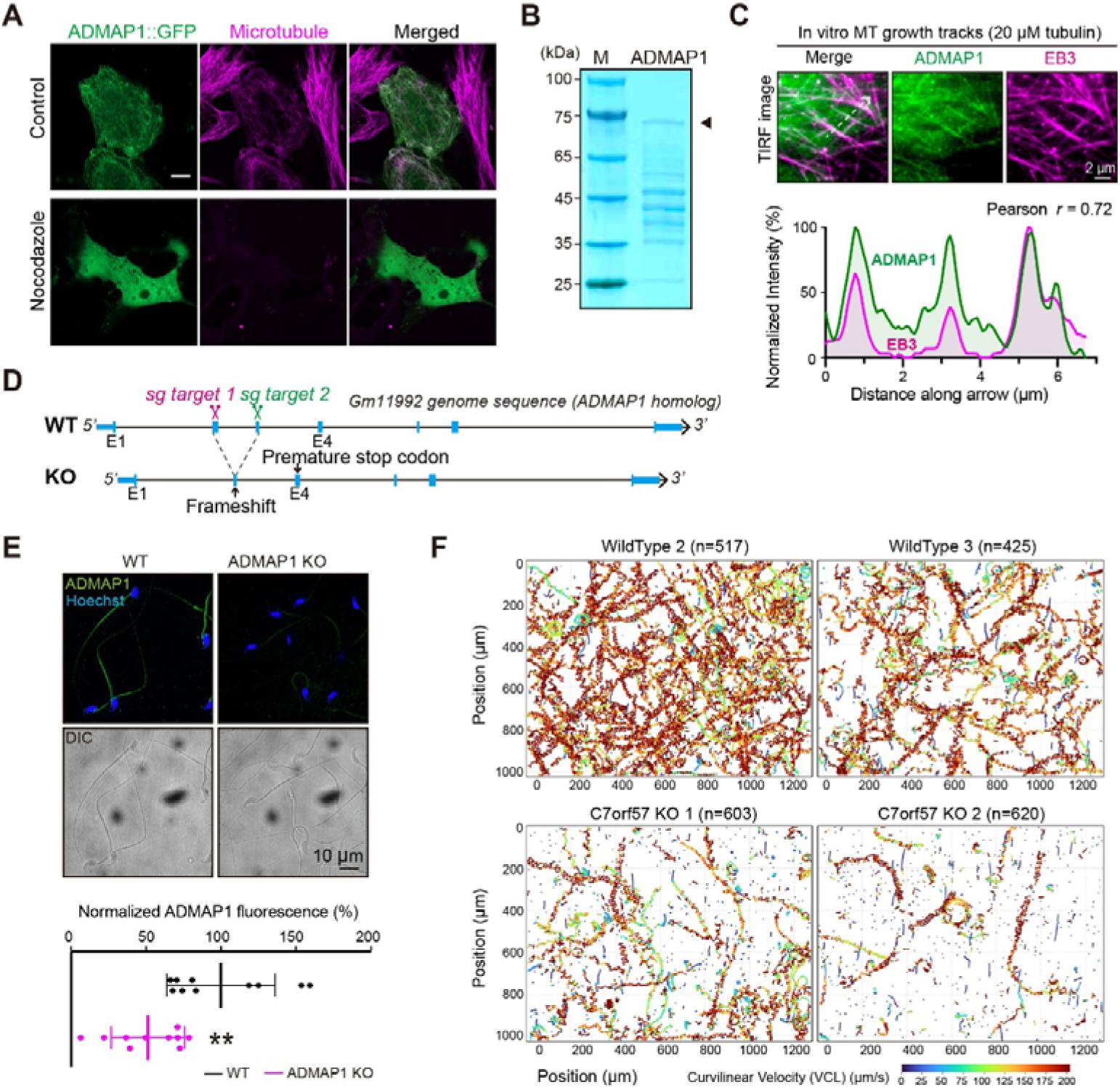
Functional and ultrastructural characterization of C7orf57. (**A**) Localization of C7orf57-EGFP in cells following nocodazole treatment. Under control conditions, C7orf57 colocalized with the microtubule cytoskeleton, whereas nocodazole-mediated microtubule depolymerization disrupted filamentous localization. Microtubules were visualized using rhodamine-labeled tubulin staining. Scale bar, 10 μm. (**B**) Coomassie-stained SDS–PAGE gel showing purified recombinant C7orf57 protein. Arrowhead indicates the major C7orf57 band. (**C**) Top: Maximum intensity projection of Z-stack TIRF images showing ADMAP1 (green) and EB3 (magenta) tracking polymerizing MT ends (20 μM tubulin). Bottom: Normalized intensity profiles along the dashed arrow (merge panel) reveal strong spatial correlation (Pearson r = 0.72) between the two proteins. Scale bar: 2 μm. (**D**) Schematic of the C7orf57 knockout strategy. CRISPR-targeted frameshift mutations introduced premature stop codons within the C7orf57 homologous locus. (**E**) Immunofluorescence staining of endogenous C7orf57 in wild-type and C7orf57 knockout mouse sperm. C7orf57 signal is shown in green and nuclei were counterstained with Hoechst (blue). Lower, quantification of normalized C7orf57 fluorescence intensity. Error bars indicate standard deviation. Statistical significance was evaluated using two-sided Mann–Whitney U tests. **P < 0.01. Scale bar, 10 μm. (**F**) Representative sperm motility trajectories from wild-type and C7orf57 knockout animals. Trajectories are colored according to curvilinear velocity (VCL). Numbers of analyzed sperm tracks are indicated above each panel.

**Figure S3.**
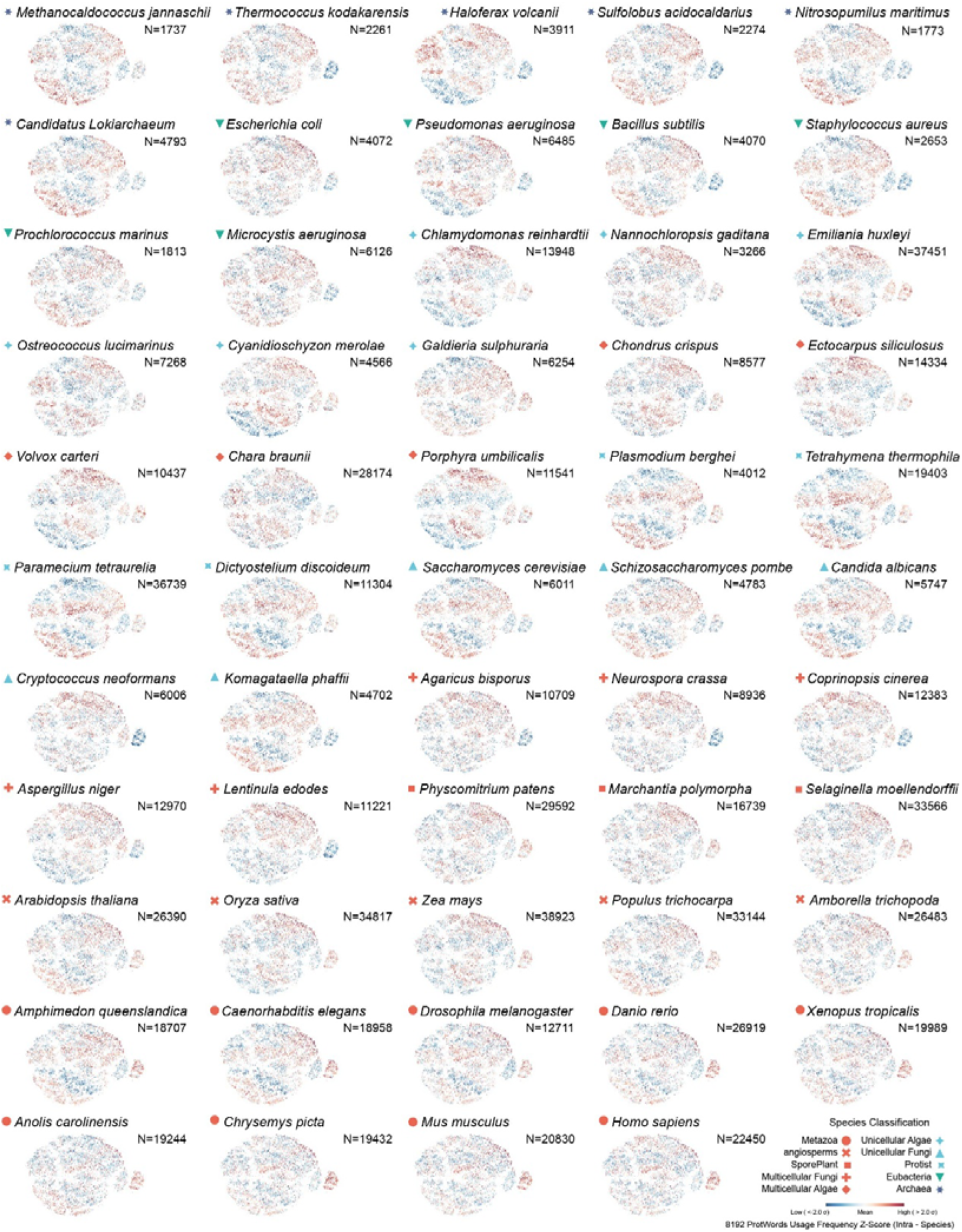
Comparative ProtWord usage landscapes reveal conserved and lineage-specific organizational patterns across the tree of life.

**Figure S4.**
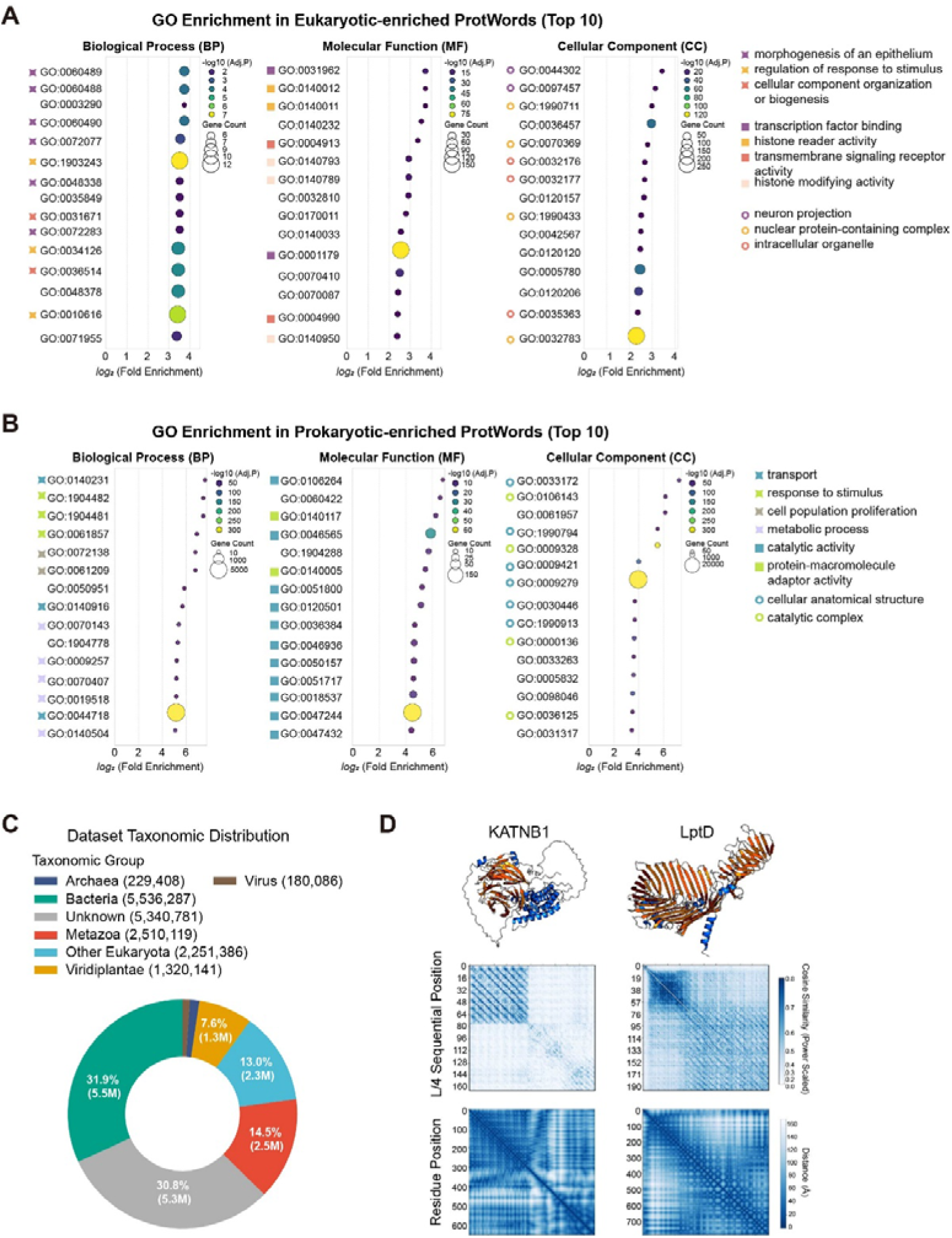
Functional and organizational properties of ProtWord latent space across evolutionary domains. (**A**) Gene Ontology (GO) enrichment analysis of the top 10 ProtWords associated with eukaryotic-enriched regions of the ICA landscape. Enriched biological process (BP), molecular function (MF), and cellular component (CC) terms are shown. (**B**) Gene Ontology (GO) enrichment analysis of the top 10 ProtWords associated with prokaryotic-enriched regions of the ICA landscape. Enriched biological process (BP), molecular function (MF), and cellular component (CC) terms are shown. (**C**) Taxonomic distribution of UniRef50 sequences used as the background dataset for GO enrichment analysis. (**D**) Additional examples comparing ProtWord latent similarity organization with protein structural organization. Top panels show representative protein structures. Middle panels show cosine similarity matrices between ProtWord latent representations across compressed sequential positions. Bottom panels show corresponding residue distance matrices.

**Figure S5.**
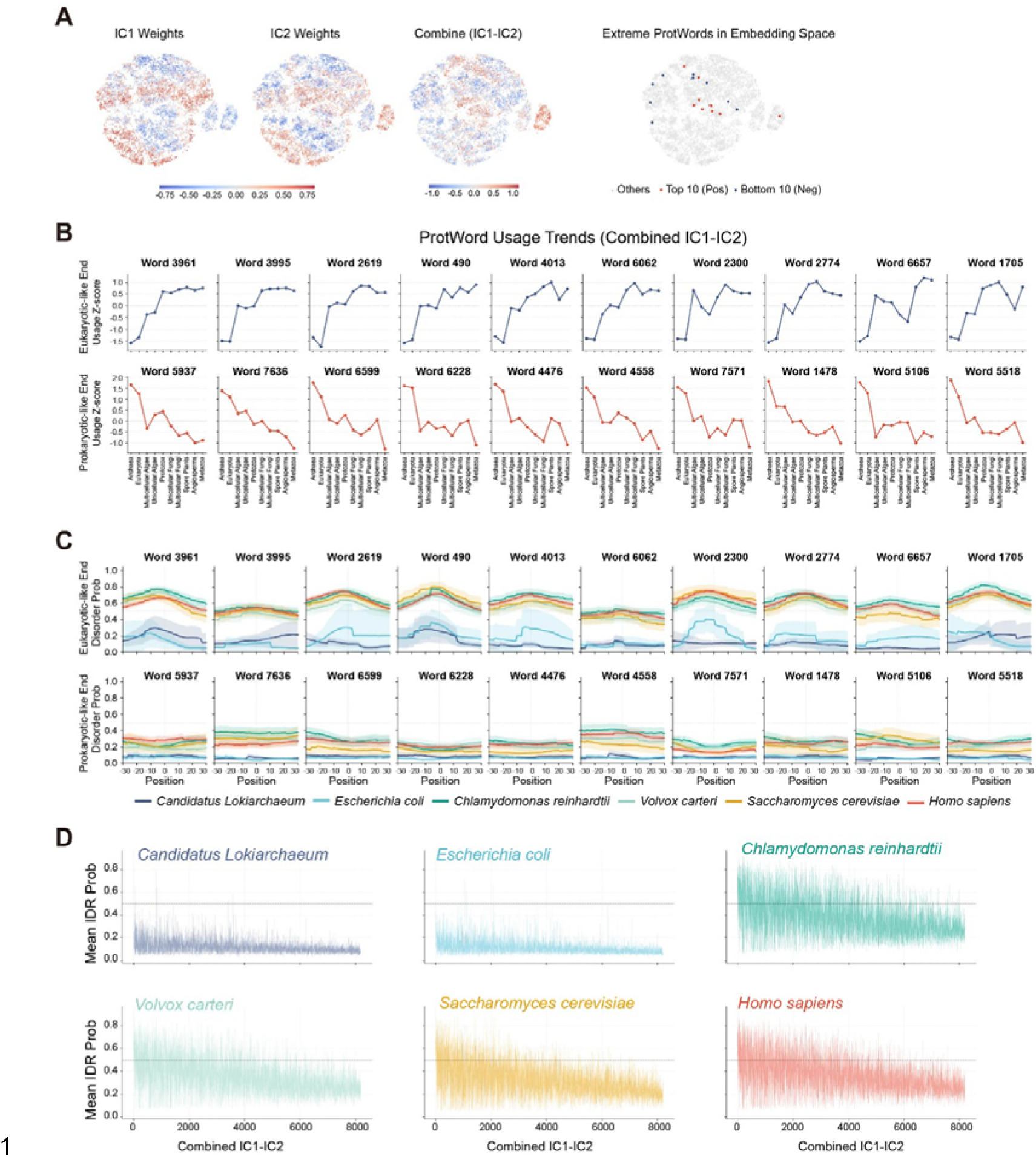
Evolutionary organization and disorder-associated trends across the ProtWord latent landscape. (**A**) Independent component analysis (ICA) of ProtWord usage representations. Left panels show ProtWord weight distributions across IC1, IC2, and the combined IC1–IC2 axis. Right panel highlights the top 10 positively weighted and bottom 10 negatively weighted ProtWords within the latent embedding landscape. (**B**) Evolutionary usage trends of representative ProtWords across major taxonomic groups ordered along the combined IC1–IC2 axis from prokaryotic to eukaryotic-enriched regions. Values represent normalized ProtWord usage Z-scores across species. (**C**) Local intrinsic disorder distributions surrounding representative ProtWords across evolutionary lineages. For each ProtWord, surrounding sequence regions were fourfold upsampled and analyzed across ±30 amino acid positions relative to the ProtWord center. Disorder probabilities were estimated from AlphaFold Database structural models sampled from representative species. (**D**) Local intrinsic disorder distributions surrounding representative ProtWords across evolutionary lineages. For each ProtWord, surrounding sequence regions were fourfold upsampled and analyzed across ±30 amino acid positions relative to the ProtWord center. Intrinsic disorder probabilities were predicted directly from amino acid sequences using MetaPredict v2 across proteins sampled from representative species.

**Figure S6.**
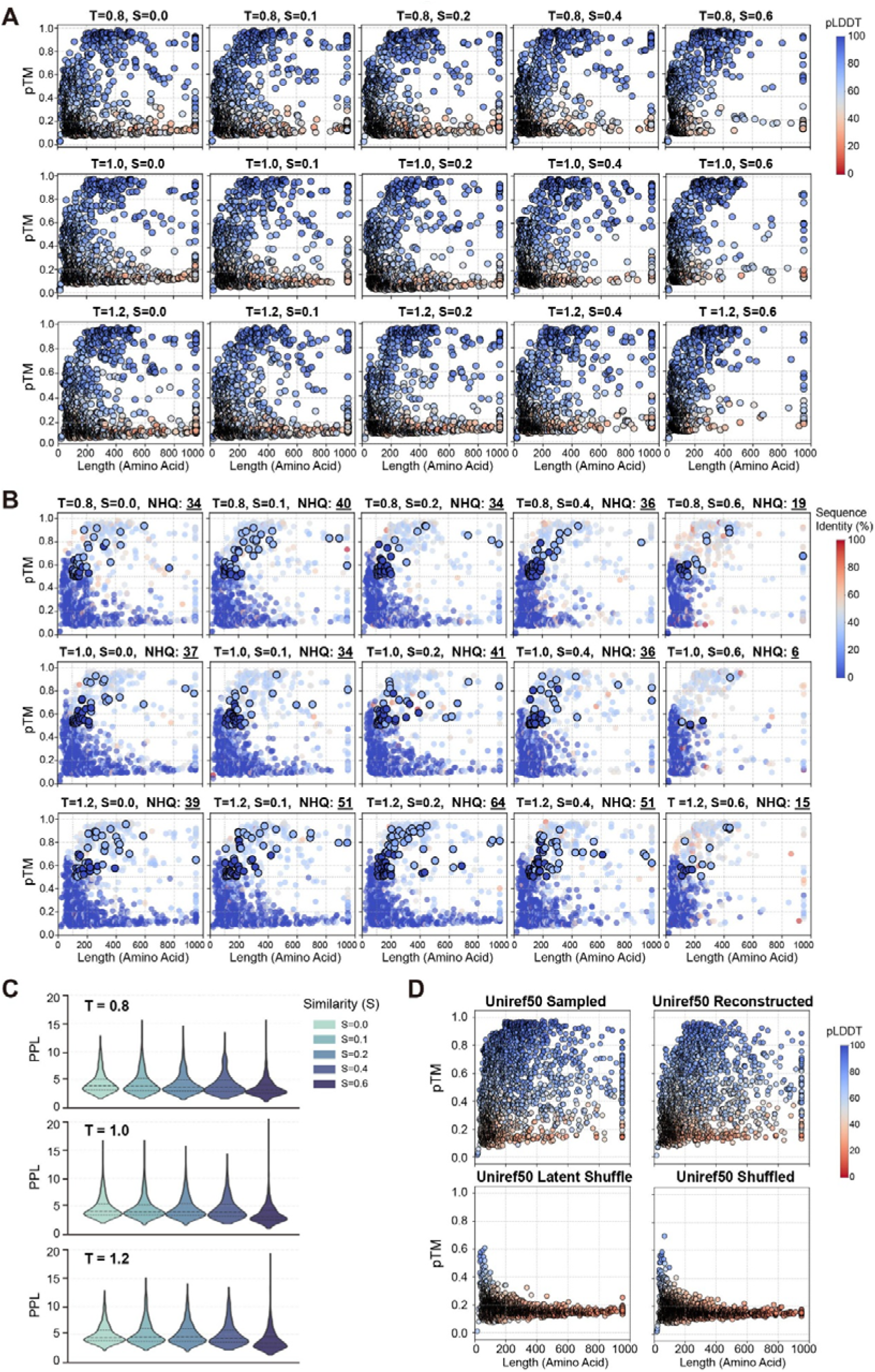
Structural quality and novelty assessment of ProtWord-based protein generation across sampling conditions. (**A**) Structural quality distributions of generated proteins across different sampling temperatures (T) and ProtWord similarity constraints (S). For each condition, 1,000 sequences were generated and evaluated using ESMFold. Predicted TM-score (pTM) is plotted against protein length, with points colored by mean pLDDT confidence scores. (**B**) Sequence novelty analysis of generated proteins under different sampling temperatures (T) and ProtWord similarity constraints (S). Predicted TM-score (pTM) is plotted against protein length, with points colored according to sequence identity to the nearest UniRef50 sequence identified using MMseqs2. Black-circled points indicate novel high-quality (NHQ) sequences, defined as proteins longer than 100 amino acids with pTM > 0.5 and sequence identity < 30% relative to the nearest UniRef50 homolog. (**C**) Perplexity distributions of generated protein sequences under different sampling temperatures (T) and ProtWord similarity constraints (S), evaluated using ESM2-650M. (**D**) Foldability assessment of UniRef50-derived sequences under reconstruction and sequence perturbation conditions. A total of 2,000 sequences randomly sampled from UniRef50 were subjected to VQ-VAE reconstruction, ProtWord-level latent shuffling, or amino acid-level shuffling, followed by structure prediction using ESMFold. Predicted TM-score (pTM) is plotted against sequence length, and points are colored according to mean predicted local distance difference test (pLDDT) confidence scores.

**Figure S7.**
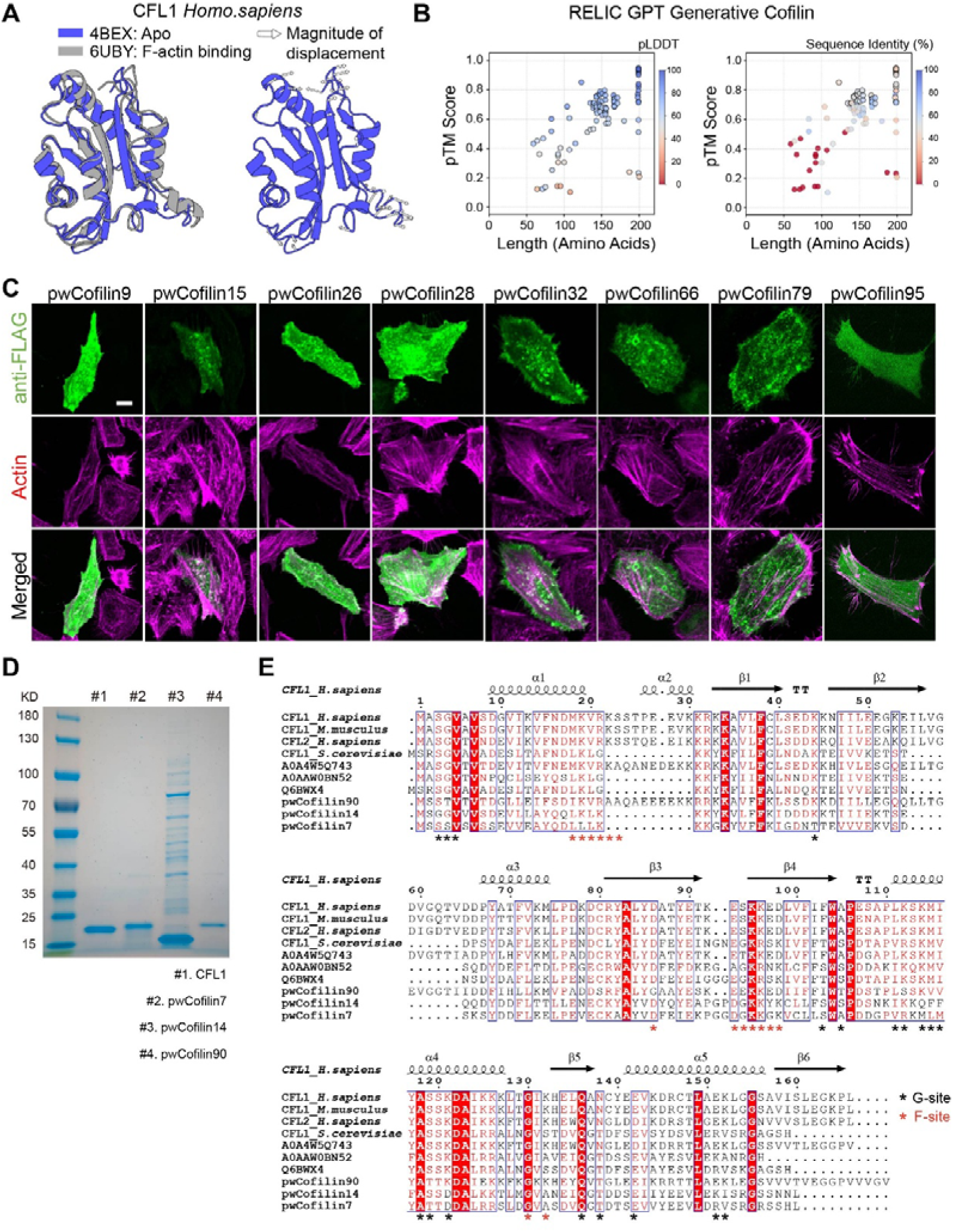
Structural comparison, sequence analysis, cellular localization, and purification of generated cofilin-like proteins. (**A**) Structural comparison of cofilin in apo and F-actin–bound conformations. Human cofilin structures in the apo state (PDB: 4BEX) and actin-bound state (PDB: 6UBY) are superimposed to illustrate conformational rearrangements associated with filament binding. Arrows indicate the predominant direction of structural displacement upon actin engagement. (**B**) Structural confidence and sequence novelty of 100 generated cofilin-like proteins predicted using ESMFold. Predicted TM-score (pTM) is plotted against protein length. In the left panel, points are colored according to mean pLDDT confidence scores. In the right panel, points are colored according to sequence identity to the nearest UniRef100 homolog. (**C**) Representative cellular localization patterns of additional generated cofilin-like proteins following transient expression in mammalian cells. Cells were stained with anti-FLAG antibody (green) and fluorescent phalloidin to visualize F-actin (magenta). Most generated proteins exhibited diffuse cytoplasmic expression without obvious aggregation, although only a subset induced strong actin filament disruption phenotypes. Scale bar, 10 μm. (**D**) Coomassie-stained SDS–PAGE analysis of purified recombinant proteins used for biochemical assays. Lane #1, human CFL1; lane #2, pwCofilin7; lane #3, pwCofilin14; lane #4, pwCofilin90. (**E**) Multiple sequence alignment of experimentally characterized generated cofilin-like proteins (pwCofilin7, pwCofilin14, and pwCofilin90), their nearest UniRef100 homologs, and representative natural cofilins from human, mouse, and yeast. Conserved residues are highlighted, and canonical cofilin functional regions, including the G-site and F-site, are indicated. Despite substantial divergence across multiple conserved positions and local structural regions, functional generated variants retained actin-remodeling activity.

