## Supplementary material for "Hierarchical latent representations reveal protein organization for functional discovery and design": Materials and Methods

**Supplementary materials**

### Materials and Methods

#### Dataset preparation

##### Pre-training Dataset (UniRef 50)

Evolutionary protein sequences were sourced from the UniRef50 database (Release 2025_04). To strictly prevent data leakage across data splits, we performed an additional clustering step on this dataset using MMseqs2 (Steinegger & Söding, 2017) at 50% sequence similarity. The resulting clusters were randomly partitioned into training (58,583,014 sequences), validation (175,616 sequences), and test (117,351 sequences) sets. This consistent cluster-based split was applied to all training stages, including the RELIC encoder, the VQ-VAE, and the latent GPT.

To efficiently manage variable sequence lengths during the training of the RELIC encoder and the VQ-VAE, a dynamic cropping strategy was applied. Sequences exceeding the context window (L=1024) were subjected to random cropping, whereas shorter sequences were padded to the required length.

Prior to training the RELIC latent GPT, sequences exceeding 1,500 residues were filtered out from the partitioned datasets to optimize computational efficiency. These filtered datasets were subsequently processed into discrete representations using the frozen weights of the pre-trained RELIC encoder and VQ-VAE.

##### Cofilin Family data for fine-tuning

A dataset of 1,611 Cofilin/ADF sequences, collected from the UniRef100 database via Pfam annotation and length filtering, was processed using rigorous homology partitioning. We employed MMseqs2 to cluster these sequences at a 50% sequence identity threshold, resulting in a training set of 1,475 sequences and a validation set of 136 sequences. This cluster-based split ensures that the validation set evaluates the model's generalization to distinct sub-families rather than mere memorization.

##### Proteome data for evolutionary analysis

All the proteomes used in evolutionary analysis and their download URLs are listed as follows:

| Species | Proteome source used in this study |
| --- | --- |
| *Methanocaldococcus jannaschii* | https://ftp.ensemblgenomes.ebi.ac.uk/pub/bacteria/release-62/fasta/bacteria_0_collection/methanocaldococcus_jannaschii_dsm_2661_gca_000091665/pep/Methanocaldococcus_jannaschii_dsm_2661_gca_000091665.ASM9166v1.pep.all.fa.gz |
| *Thermococcus kodakarensis* | https://ftp.ensemblgenomes.ebi.ac.uk/pub/bacteria/release-62/fasta/bacteria_0_collection/thermococcus_kodakarensis_kod1_gca_000009965/pep/Thermococcus_kodakarensis_kod1_gca_000009965.ASM996v1.pep.all.fa.gz |
| *Haloferax volcanii* | https://ftp.ensemblgenomes.ebi.ac.uk/pub/bacteria/release-62/fasta/bacteria_0_collection/haloferax_volcanii_ds2_gca_000025685/pep/Haloferax_volcanii_ds2_gca_000025685.ASM2568v1.pep.all.fa.gz |
| *Sulfolobus acidocaldarius* | https://ftp.ensemblgenomes.ebi.ac.uk/pub/bacteria/release-62/fasta/bacteria_126_collection/sulfolobus_acidocaldarius_dsm_639_gca_000012285/pep/Sulfolobus_acidocaldarius_dsm_639_gca_000012285.ASM1228v1.pep.all.fa.gz |
| *Nitrosopumilus maritimus* | https://ftp.ensemblgenomes.ebi.ac.uk/pub/bacteria/release-62/fasta/bacteria_0_collection/nitrosopumilus_maritimus_scm1_gca_000018465/pep/Nitrosopumilus_maritimus_scm1_gca_000018465.ASM1846v1.pep.all.fa.gz |
| *Candidatus Lokiarchaeum* | https://ftp.ensemblgenomes.ebi.ac.uk/pub/bacteria/release-62/fasta/bacteria_3_collection/candidatus_lokiarchaeum_sp_gc14_75_gca_000986845/pep/Candidatus_lokiarchaeum_sp_gc14_75_gca_000986845.ASM98684v1.pep.all.fa.gz |
| *Escherichia coli* | https://ftp.ensemblgenomes.ebi.ac.uk/pub/bacteria/release-62/fasta/bacteria_0_collection/escherichia_coli_str_k_12_substr_mg1655_gca_000005845/pep/Escherichia_coli_str_k_12_substr_mg1655_gca_000005845.ASM584v2.pep.all.fa.gz |
| *Pseudomonas aeruginosa* | https://ftp.ensemblgenomes.ebi.ac.uk/pub/bacteria/release-62/fasta/bacteria_3_collection/pseudomonas_aeruginosa_gca_001051345/pep/Pseudomonas_aeruginosa_gca_001051345.PAMH19.pep.all.fa.gz |
| *Bacillus subtilis* | https://ftp.ensemblgenomes.ebi.ac.uk/pub/bacteria/release-62/fasta/bacteria_0_collection/bacillus_subtilis_subsp_subtilis_str_168_gca_000009045/pep/Bacillus_subtilis_subsp_subtilis_str_168_gca_000009045.ASM904v1.pep.all.fa.gz |
| *Staphylococcus aureus* | https://ftp.ensemblgenomes.ebi.ac.uk/pub/bacteria/release-62/fasta/bacteria_2_collection/staphylococcus_aureus_gca_001049575/pep/Staphylococcus_aureus_gca_001049575.AH1.pep.all.fa.gz |
| *Prochlorococcus marinus* | https://ftp.ensemblgenomes.ebi.ac.uk/pub/bacteria/release-62/fasta/bacteria_0_collection/prochlorococcus_marinus_subsp_marinus_str_ccmp1375_gca_000007925/pep/Prochlorococcus_marinus_subsp_marinus_str_ccmp1375_gca_000007925.ASM792v1.pep.all.fa.gz |
| *Microcystis aeruginosa* | https://ftp.ensemblgenomes.ebi.ac.uk/pub/bacteria/release-62/fasta/bacteria_0_collection/microcystis_aeruginosa_nies_843_gca_000010625/pep/Microcystis_aeruginosa_nies_843_gca_000010625.ASM1062v1.pep.all.fa.gz |
| *Amphimedon queenslandica* | https://ftp.ensemblgenomes.ebi.ac.uk/pub/metazoa/release-62/fasta//amphimedon_queenslandica_gca000090795v2rs/pep/Amphimedon_queenslandica_gca000090795v2rs.v1.1.pep.all.fa.gz |
| *Caenorhabditis elegans* | https://ftp.ensembl.org/pub/release-115/fasta/caenorhabditis_elegans/pep/Caenorhabditis_elegans.WBcel235.pep.all.fa.gz |
| *Drosophila melanogaster* | https://ftp.ensembl.org/pub/release-115/fasta/drosophila_melanogaster/pep/Drosophila_melanogaster.BDGP6.54.pep.all.fa.gz |
| *Danio rerio* | https://ftp.ensembl.org/pub/release-115/fasta/danio_rerio/pep/Danio_rerio.GRCz11.pep.all.fa.gz |
| *Xenopus tropicalis* | https://ftp.ensembl.org/pub/release-115/fasta/xenopus_tropicalis/pep/Xenopus_tropicalis.UCB_Xtro_10.0.pep.all.fa.gz |
| *Anolis carolinensis* | https://ftp.ensembl.org/pub/release-115/fasta/anolis_carolinensis/pep/Anolis_carolinensis.AnoCar2.0v2.pep.all.fa.gz |
| *Chrysemys picta* | https://ftp.ensembl.org/pub/release-115/fasta/chrysemys_picta_bellii/pep/Chrysemys_picta_bellii.Chrysemys_picta_bellii-3.0.3.pep.all.fa.gz |
| *Mus musculus* | https://ftp.ensembl.org/pub/release-115/fasta/mus_musculus/pep/Mus_musculus.GRCm39.pep.all.fa.gz |
| *Homo sapiens* | https://ftp.ensembl.org/pub/release-115/fasta/homo_sapiens/pep/Homo_sapiens.GRCh38.pep.all.fa.gz |
| *Chondrus crispus* | https://ftp.ensemblgenomes.ebi.ac.uk/pub/plants/release-62/fasta//chondrus_crispus/pep/Chondrus_crispus.ASM35022v2.pep.all.fa.gz |
| *Ectocarpus siliculosus* | https://ftp.ensemblgenomes.ebi.ac.uk/pub/protists/release-62/fasta/protists_stramenopiles1_collection/ectocarpus_siliculosus_gca_000310025/pep/Ectocarpus_siliculosus_gca_000310025.ASM31002v1.pep.all.fa.gz |
| *Volvox carteri* | https://ftp.ncbi.nlm.nih.gov/genomes/all/GCF/000/143/455/GCF_000143455.1_v1.0/GCF_000143455.1_v1.0_protein.faa.gz |
| *Chara braunii* | https://ftp.ensemblgenomes.ebi.ac.uk/pub/plants/release-62/fasta//chara_braunii/pep/Chara_braunii.Cbr_1.0.pep.all.fa.gz |
| *Porphyra umbilicalis* | https://ftp.ncbi.nlm.nih.gov/genomes/all/GCA/002/049/455/GCA_002049455.2_P_umbilicalis_v1/GCA_002049455.2_P_umbilicalis_v1_protein.faa.gz |
| *Agaricus bisporus* | https://ftp.ensemblgenomes.ebi.ac.uk/pub/fungi/release-62/fasta/fungi_basidiomycota1_collection/agaricus_bisporus_var_burnettii_jb137_s8_gca_000300555/pep/Agaricus_bisporus_var_burnettii_jb137_s8_gca_000300555.Agabi_varbur_1.pep.all.fa.gz |
| *Neurospora crassa* | https://ftp.ensemblgenomes.ebi.ac.uk/pub/fungi/release-62/fasta//neurospora_crassa/pep/Neurospora_crassa.NC12.pep.all.fa.gz |
| *Coprinopsis cinerea* | https://ftp.ensemblgenomes.ebi.ac.uk/pub/fungi/release-62/fasta/fungi_basidiomycota1_collection/coprinopsis_cinerea_okayama7_130_gca_000182895/pep/Coprinopsis_cinerea_okayama7_130_gca_000182895.CC3.pep.all.fa.gz |
| *Aspergillus niger* | https://ftp.ensemblgenomes.ebi.ac.uk/pub/fungi/release-62/fasta//aspergillus_niger/pep/Aspergillus_niger.ASM285v2.pep.all.fa.gz |
| *Lentinula edodes* | https://ftp.ensemblgenomes.ebi.ac.uk/pub/fungi/release-62/fasta/fungi_basidiomycota1_collection/lentinula_edodes_gca_002003045/pep/Lentinula_edodes_gca_002003045.Lened_assembly01.pep.all.fa.gz |
| *Plasmodium berghei* | https://ftp.ensemblgenomes.ebi.ac.uk/pub/protists/release-62/fasta//plasmodium_berghei/pep/Plasmodium_berghei.GCA900002375v2.pep.all.fa.gz |
| *Tetrahymena thermophila* | https://ftp.ensemblgenomes.ebi.ac.uk/pub/protists/release-62/fasta//tetrahymena_thermophila/pep/Tetrahymena_thermophila.JCVI-TTA1-2.2.pep.all.fa.gz |
| *Paramecium tetraurelia* | https://ftp.ensemblgenomes.ebi.ac.uk/pub/protists/release-62/fasta//paramecium_tetraurelia/pep/Paramecium_tetraurelia.ASM16542v1.pep.all.fa.gz |
| *Dictyostelium discoideum* | https://ftp.ensemblgenomes.ebi.ac.uk/pub/protists/release-62/fasta/dictyostelium_discoideum/pep/Dictyostelium_discoideum.dicty_2.7.pep.all.fa.gz |
| *Physcomitrium patens* | https://ftp.ensemblgenomes.ebi.ac.uk/pub/plants/release-62/fasta//physcomitrium_patens/pep/Physcomitrium_patens.Phypa_V3.pep.all.fa.gz |
| *Marchantia polymorpha* | https://ftp.ensemblgenomes.ebi.ac.uk/pub/plants/release-62/fasta//marchantia_polymorpha/pep/Marchantia_polymorpha.MpTak1_v7.1.pep.all.fa.gz |
| *Selaginella moellendorffii* | https://ftp.ensemblgenomes.ebi.ac.uk/pub/plants/release-62/fasta//selaginella_moellendorffii/pep/Selaginella_moellendorffii.v1.0.pep.all.fa.gz |
| *Chlamydomonas reinhardtii* | https://ftp.ensemblgenomes.ebi.ac.uk/pub/plants/release-62/fasta//chlamydomonas_reinhardtii/pep/Chlamydomonas_reinhardtii.Chlamydomonas_reinhardtii_v5.5.pep.all.fa.gz |
| *Nannochloropsis gaditana* | https://ftp.ensemblgenomes.ebi.ac.uk/pub/protists/release-62/fasta/protists_stramenopiles1_collection/nannochloropsis_gaditana_ccmp526_gca_000240725/pep/Nannochloropsis_gaditana_ccmp526_gca_000240725.ASM24072v1.pep.all.fa.gz |
| *Emiliania huxleyi* | https://ftp.ensemblgenomes.ebi.ac.uk/pub/protists/release-62/fasta//emiliania_huxleyi/pep/Emiliania_huxleyi.Emiliana_huxleyi_CCMP1516_main_genome_assembly_v1.0.pep.all.fa.gz |
| *Ostreococcus lucimarinus* | https://ftp.ensemblgenomes.ebi.ac.uk/pub/plants/release-62/fasta//ostreococcus_lucimarinus/pep/Ostreococcus_lucimarinus.ASM9206v1.pep.all.fa.gz |
| *Cyanidioschyzon merolae* | https://ftp.ensemblgenomes.ebi.ac.uk/pub/plants/release-62/fasta//cyanidioschyzon_merolae/pep/Cyanidioschyzon_merolae.ASM9120v1.pep.all.fa.gz |
| *Galdieria sulphuraria* | https://ftp.ensemblgenomes.ebi.ac.uk/pub/plants/release-62/fasta//galdieria_sulphuraria/pep/Galdieria_sulphuraria.ASM34128v1.pep.all.fa.gz |
| *Saccharomyces cerevisiae* | https://ftp.ensembl.org/pub/release-115/fasta/saccharomyces_cerevisiae/pep/Saccharomyces_cerevisiae.R64-1-1.pep.all.fa.gz |
| *Schizosaccharomyces pombe* | https://ftp.ensemblgenomes.ebi.ac.uk/pub/fungi/release-62/fasta//schizosaccharomyces_pombe/pep/Schizosaccharomyces_pombe.ASM294v2.pep.all.fa.gz |
| *Candida albicans* | https://ftp.ensemblgenomes.ebi.ac.uk/pub/fungi/release-62/fasta//candida_albicans/pep/Candida_albicans.GCA000182965v3.pep.all.fa.gz |
| *Cryptococcus neoformans* | https://ftp.ensemblgenomes.ebi.ac.uk/pub/fungi/release-62/fasta//cryptococcus_neoformans/pep/Cryptococcus_neoformans.ASM9104v1.pep.all.fa.gz |
| *Komagataella phaffii* | https://ftp.ensemblgenomes.ebi.ac.uk/pub/fungi/release-62/fasta/fungi_ascomycota3_collection/komagataella_phaffii_gs115_gca_001746955/pep/Komagataella_phaffii_gs115_gca_001746955.ASM174695v1.pep.all.fa.gz |
| *Arabidopsis thaliana* | https://ftp.ensemblgenomes.ebi.ac.uk/pub/plants/release-62/fasta/arabidopsis_thaliana/pep/Arabidopsis_thaliana.TAIR10.pep.all.fa.gz |
| *Oryza sativa* | https://ftp.ensemblgenomes.ebi.ac.uk/pub/plants/release-62/fasta/oryza_sativa/pep/Oryza_sativa.IRGSP-1.0.pep.all.fa.gz |
| *Zea mays* | https://ftp.ensemblgenomes.ebi.ac.uk/pub/plants/release-62/fasta//zea_mays/pep/Zea_mays.Zm-B73-REFERENCE-NAM-5.0.pep.all.fa.gz |
| *Populus trichocarpa* | https://ftp.ensemblgenomes.ebi.ac.uk/pub/plants/release-62/fasta//populus_trichocarpa/pep/Populus_trichocarpa.Pop_tri_v4.pep.all.fa.gz |
| *Amborella trichopoda* | https://ftp.ensemblgenomes.ebi.ac.uk/pub/plants/release-62/fasta//amborella_trichopoda/pep/Amborella_trichopoda.AMTR1.0.pep.all.fa.gz |

##### Gene ontology annotation for enrichment analysis

The GO annotation file containing all GO annotations for proteins in the UniProt Knowledgebase (UniProtKB) was obtained from https://ftp.ebi.ac.uk/pub/databases/GO/goa/UNIPROT/, using the GO version http://purl.obolibrary.org/obo/go/releases/2022-11-12/extensions/go-plus.owl generated on 2022-11-17 13:42. Species background statistics were calculated using the sequences recorded in this file that possessed a taxonomy assignment.

##### SCOPe dataset and structural homologue benchmark

The SCOPe40 dataset consists of single domains with experimentally determined structures. We generated this dataset by clustering SCOPe 2.01 at a 0.4 sequence identity threshold, yielding 11,211 non-redundant protein domain structures (Chandonia et al., 2022). We utilized all 11,211 structures for t-SNE dimensional reduction to visualize the latent space encoded by the RELIC encoder.

For the detailed benchmark against other searching tools, we utilized a designated SCOPe40-test dataset comprising 2,207 sequences. We deliberately chose the same test data employed in the evaluations of PLMSearch (Liu et al., 2024) and Foldseek (van Kempen et al., 2024). This approach is critical to prevent data leakage and ensure a fair comparison, as tools such as SS-predictor and MMseqs2 were trained directly on other partitions of this dataset. Furthermore, this specific split maximizes the diversity of the test set. All subsequent data analyses followed the pipeline established by PLMSearch (https://github.com/maovshao/PLMSearch/blob/main/main.ipynb).

##### Training objective

The RELIC encoder was pre-trained via a Masked Language Modeling (MLM) objective. Following the standard BERT protocol (Devlin et al., 2019), 15% of the residues in each sequence were randomly targeted for supervision: 80% were replaced by a specialized <mask> token, 10% were substituted with a random amino acid, and the remaining 10% were left unchanged. This configuration forces the model to integrate local physicochemical constraints with global structural dependencies to accurately reconstruct corrupted sequences.

Subsequently, the VQ-VAE was trained to reconstruct the original input from its discretized latent representation. During this stage, the weights of the RELIC encoder were strictly frozen to rely on the previously learned continuous representations.

Finally, the Latent GPT was trained on sequences tokenized into discrete ProtWords. By modeling the latent dependencies between these ProtWords, the GPT learns to capture the context-dependent structural consistencies inherent in the protein sequence space.

#### Model architecture

##### RELIC encoder

To explicitly model the hierarchical nature of protein folding, we developed a hybrid U-Net-inspired architecture (Ronneberger et al., 2015). Unlike standard Transformers that apply global attention across every residue, our model efficiently integrates Convolutional Neural Networks (CNNs) with self-attention (Algorithm 1):

- **Local Feature Extraction (Encoder):** Input sequences are processed by a two-layer 1D convolutional encoder with kernel sizes of 5 and 7. This design enforces a strong inductive bias for capturing local secondary structure motifs and reduces the sequence length by a factor of four via max-pooling.
- **Global Context Modeling (Bottleneck):** The compressed latent representation is processed by a Transformer bottleneck equipped with Rotary Position Embedding (RoPE) (Su et al., 2024). By operating at a reduced spatial resolution, the attention mechanism efficiently captures long-range tertiary interactions with significantly lower computational complexity.
- **Structure Reconstruction (Decoder):** Global features are upsampled via transposed convolutions and fused with local features from the encoder. Unlike standard U-Net concatenation, we employed element-wise addition for these residual skip connections to maintain computational efficiency. This restores the sequence to its original resolution for masked token prediction.

The final RELIC encoder architecture is constructed by sequentially cascading 12 complete U-Net modules. Within the bottleneck of each module, the Transformer is configured with a hidden dimension of 560 and 20 attention heads, carefully balancing representational capacity with computational efficiency.

- **Optimization:** Training was implemented in PyTorch and optimized using Mixed Precision (BFloat16) to maximize throughput. We used the AdamW optimizer with β1=0.9, β2=0.98, and a weight decay of 0.01. The learning rate followed a linear warmup schedule for the first 5,000 steps, peaking at 4e-4, followed by a linear decay. Training was conducted on a high-performance computing cluster utilizing 8* NVIDIA RTX4090 GPUs. Distributed training was managed via Distributed Data Parallel, with an effective batch size of 1024 sequences. The final model checkpoint used for all downstream tasks and analyses was trained for a total of 500,000 steps. (Algorithm 2)

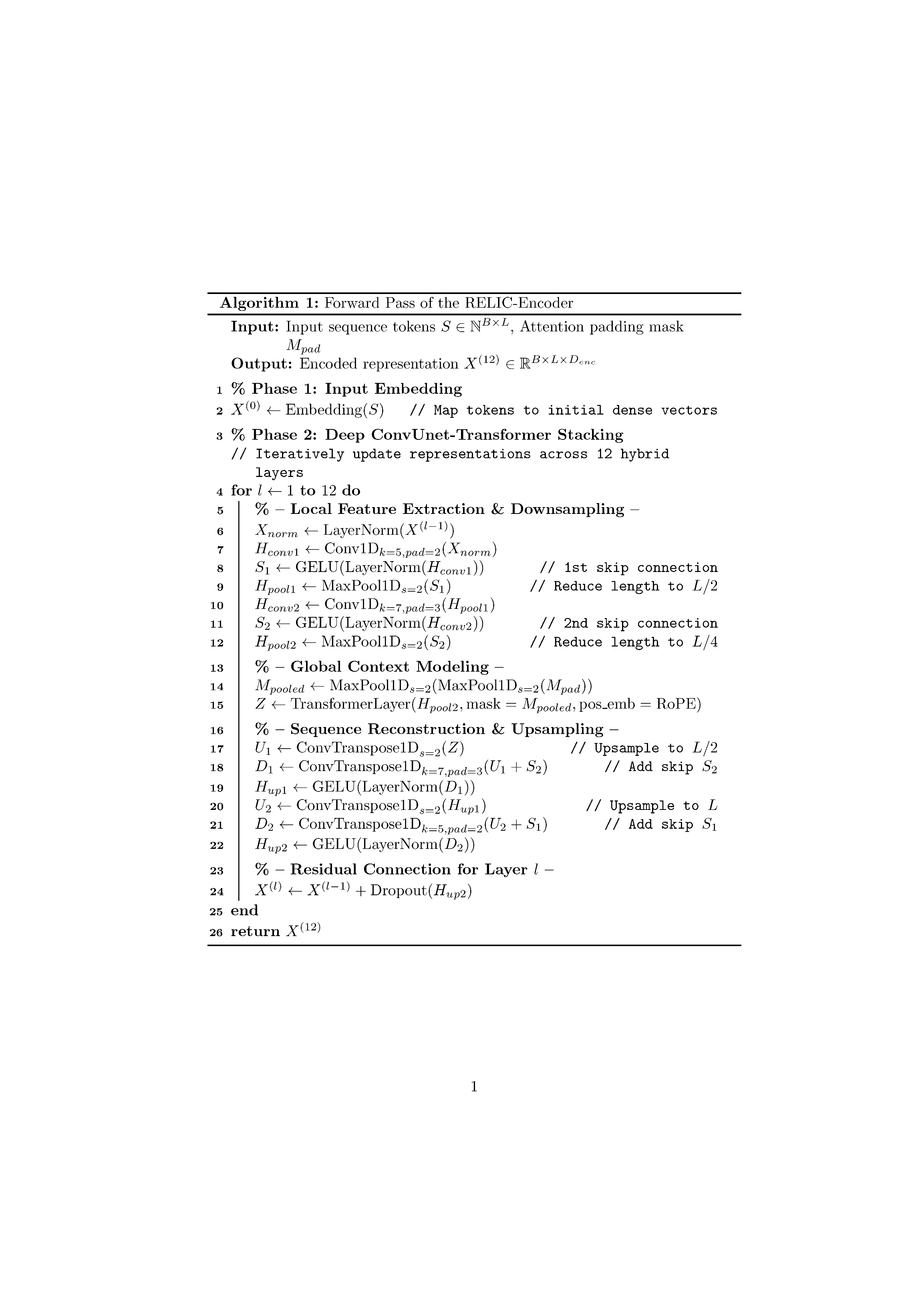

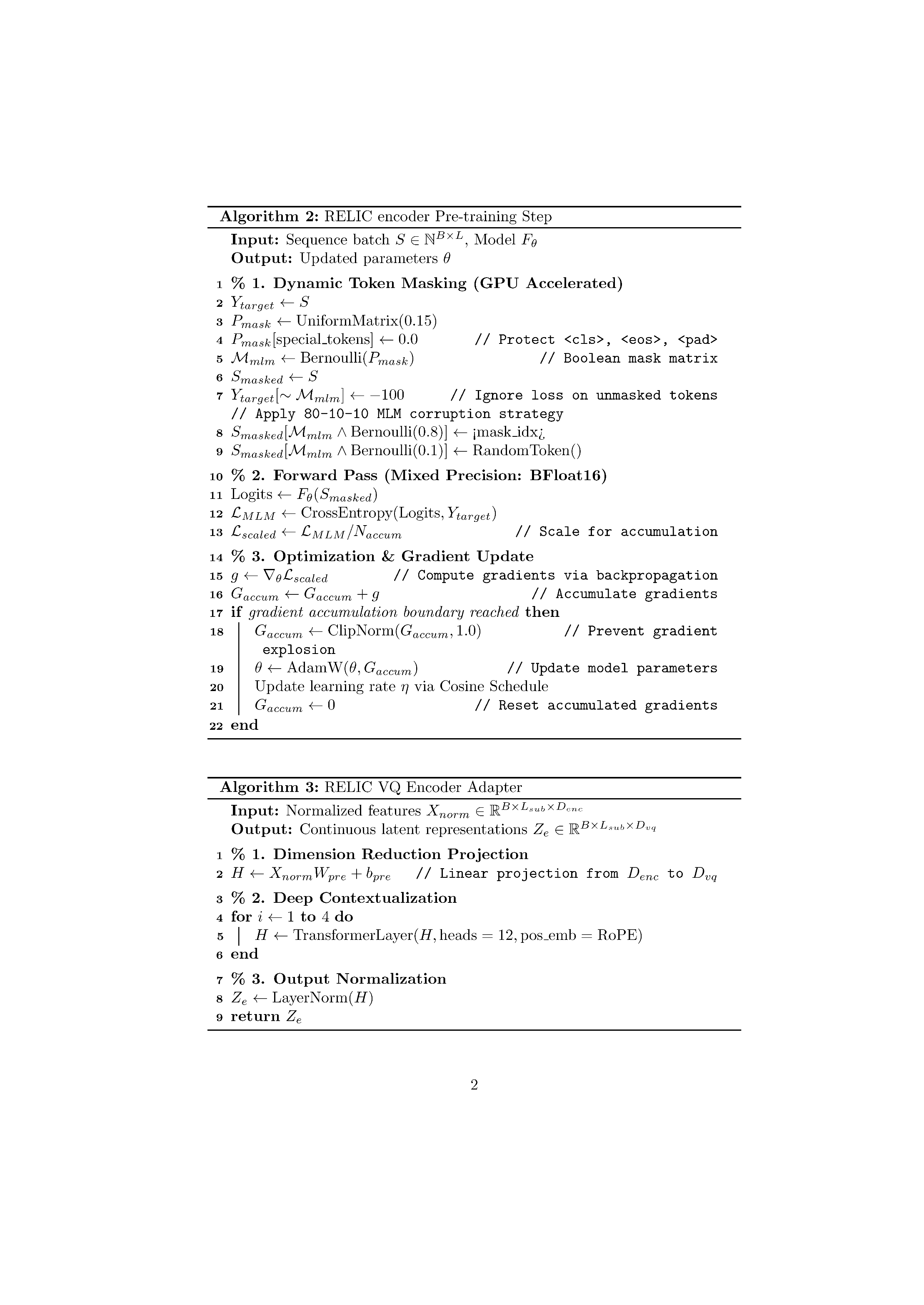

##### RELIC VQ-VAE

To transition from continuous representations to a discrete evolutionary vocabulary, we trained a Vector Quantized Variational Autoencoder (VQ-VAE) (Van Den Oord & Vinyals, 2017) on the latent features produced by the pre-trained encoder.

- **Encoder and Quantization Bottleneck:** The pre-trained hybrid encoder was utilized as a fixed feature extractor, with all weights frozen to preserve the learned topological semantics. The continuous output features were normalized and projected into a lower-dimensional manifold (d=384) via a 4-layer Transformer adapter. These features were subsequently discretized using a learnable codebook containing K=8,192 latent embeddings. To maximize codebook utilization and prevent index collapse, we employed an Exponential Moving Average (EMA) update strategy with the decay rate strictly maintained at 0.99, coupled with a dead-code revival mechanism (Algorithm 3-6).
- **Latent Topology Adaptation:** Before sequence reconstruction, it is crucial to transition the compressed, discrete tokens back into a continuous structural manifold. To bridge this gap, the quantized embeddings (d=384) are linearly projected back to the original dimension (d=560) and processed by a deep, 16-layer Transformer decoder adapter. This module ensures the robust recovery of continuous global dependencies prior to spatial upsampling.
- **Hierarchical Hybrid Decoder:** To effectively reverse the downsampling operations and reconstruct the fine-grained protein sequences, the adapted continuous features are fed into a hierarchical hybrid decoder. This decoder interleaves global context modeling with resolution recovery, consisting of two stages of 8-layer Transformer blocks separated by transposed convolutional layers (stride=2). This architecture progressively upsamples the spatial resolution from the latent space (L/4) back to the original sequence length (L), explicitly recovering local residue identities while maintaining structural constraints.
- **Stochastic Latent Mixing:** To stabilize the non-differentiable quantization bottleneck and prevent early-stage codebook neglect, we implemented a batch-wise stochastic curriculum. Let $z_{e}$ denote the continuous features from the encoder adapter and $z_{q}$ denote the corresponding quantized embedding. During training, the input to the decoder adapter, $z_{in}$, is selected via a Bernoulli process:

$$z_{in}=\left\{ \begin{aligned} z_{e} with probability p \\ z_{q} with probability 1-p \end{aligned} \right.$$

Unlike standard linear warmup, we annealed the mixing probability $p$ non-linearly from 1.0 to 0.0 over the initial 15,000 steps using a polynomial decay schedule (power=0.2). This hard-switching strategy forces the model to incrementally rely on the discrete codebook, completely resolving the "cold-start" collapse problem while allowing the continuous latent manifold to organize effectively in early stages.

- **Optimization**: The VQ-VAE model was trained for 500,000 steps using the AdamW optimizer (learning rate of 5e-5) with an effective batch size of 512. The final objective function comprised two components: a cross-entropy loss for sequence reconstruction and a standard VQ commitment loss.

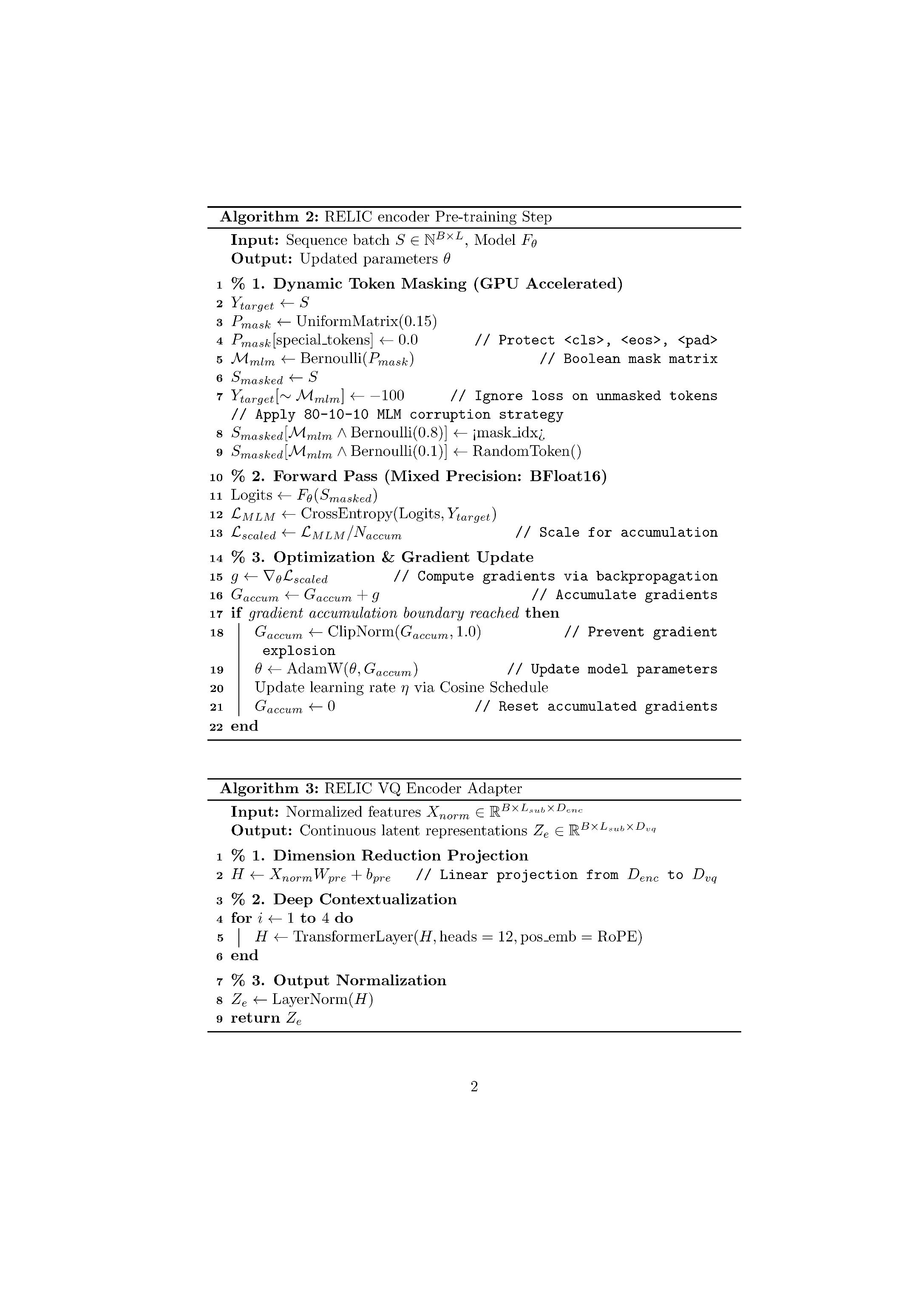

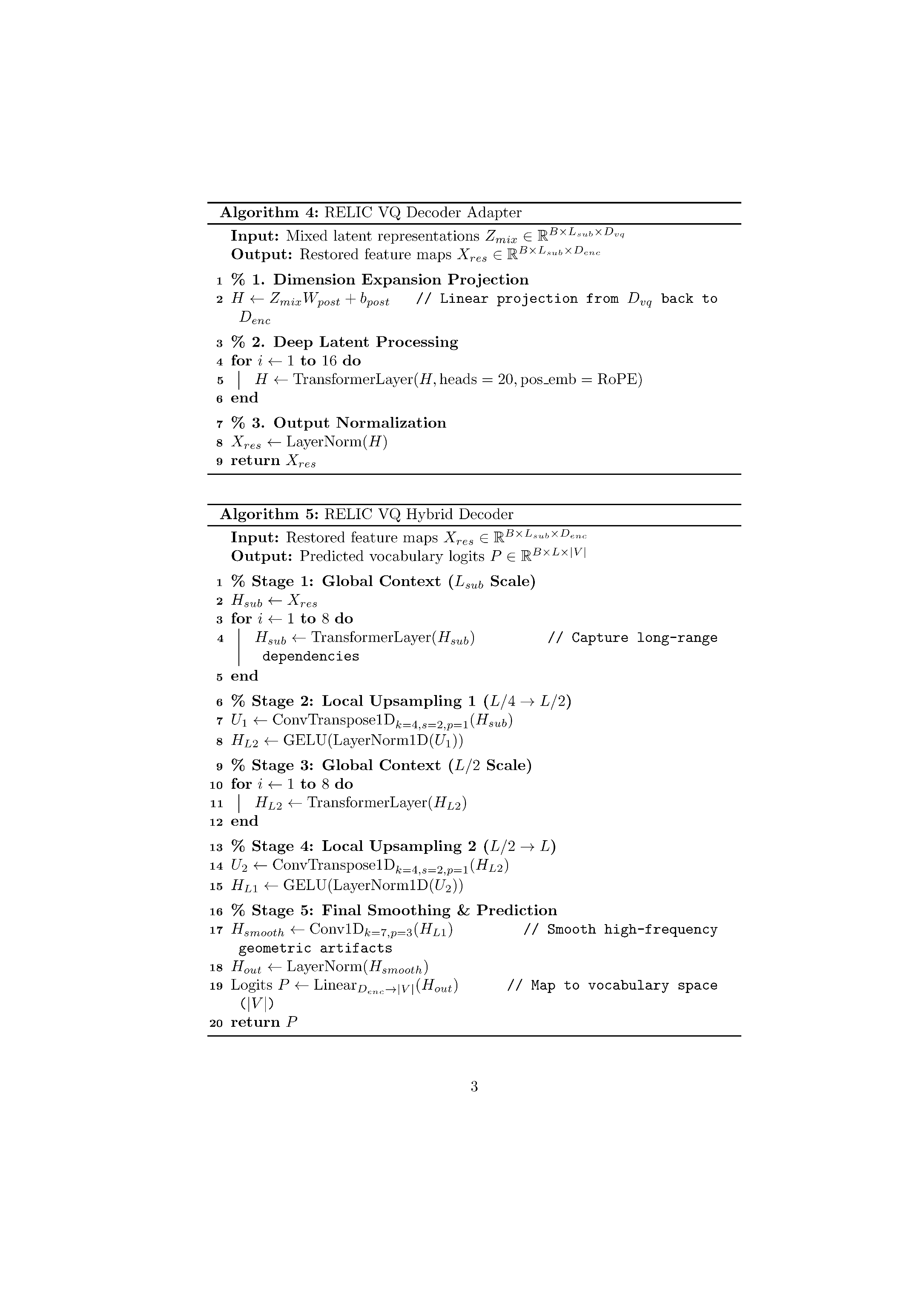

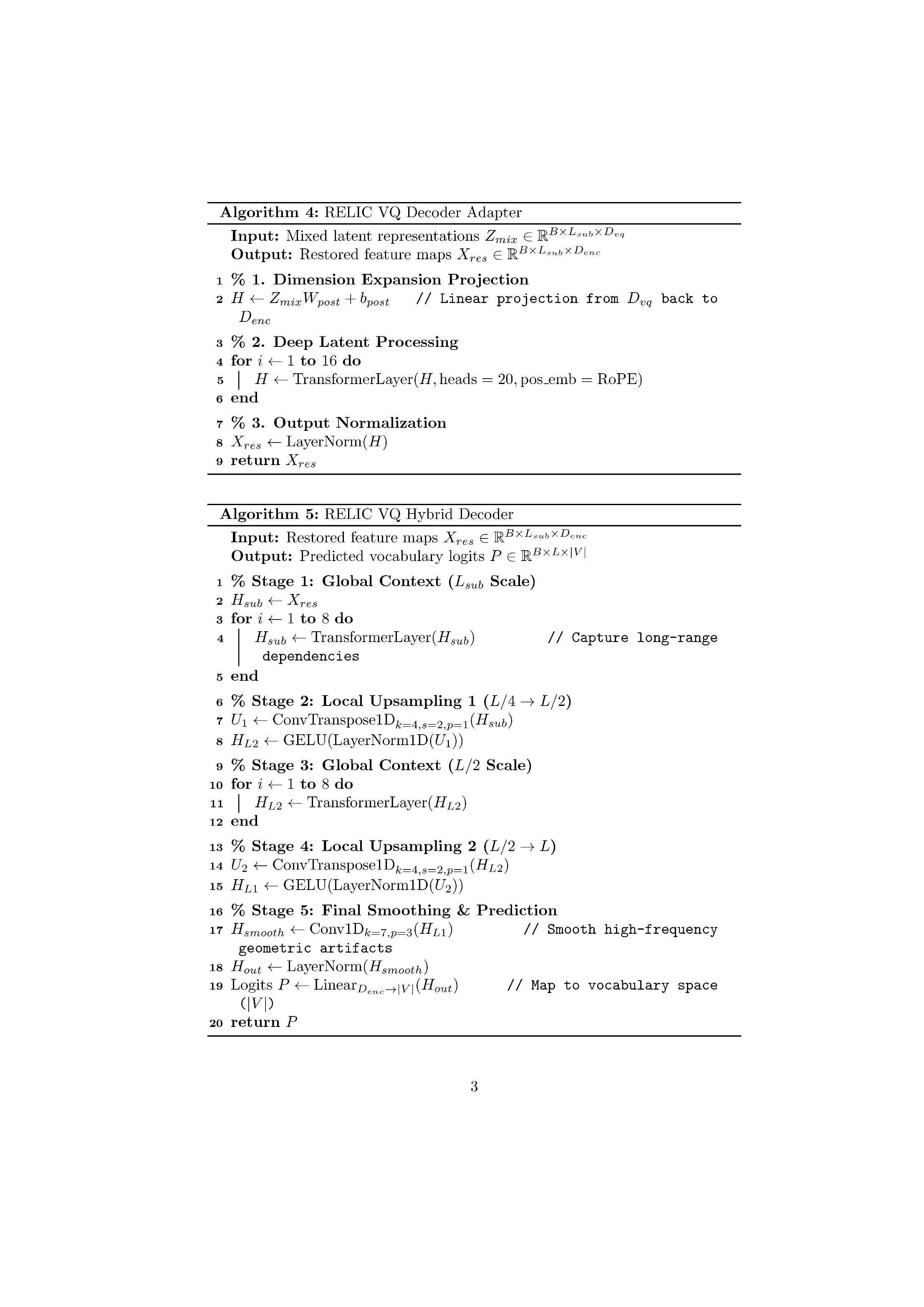

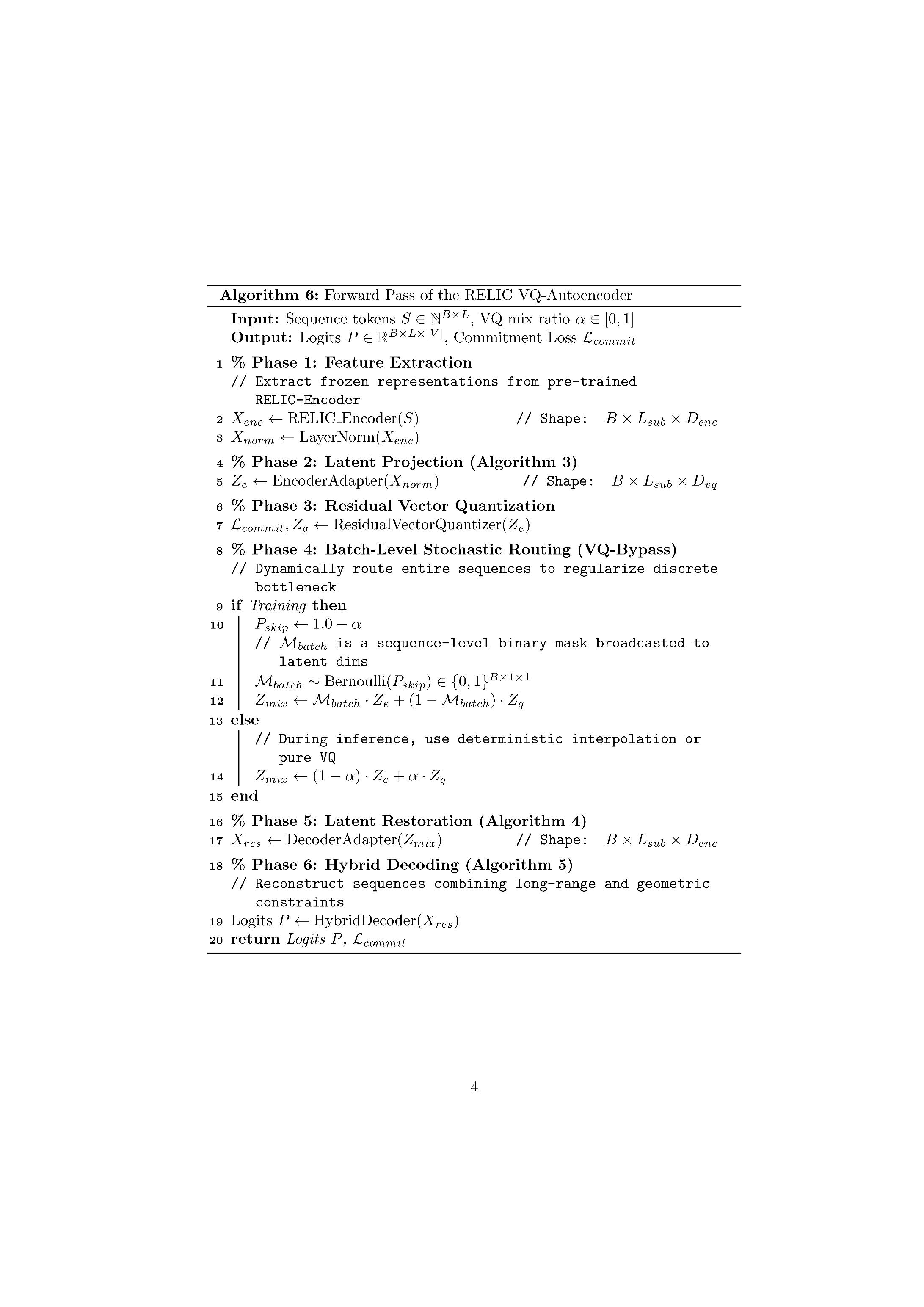

##### Latent GPT

To capture the high-level syntax and long-range dependencies of the discretized protein vocabulary, we trained a decoder-only Transformer (GPT).

The model consists of 24 layers with an embedding dimension of 768 and 12 attention heads. To maximize training stability at this depth, we incorporated several architectural enhancements （Algorithm 7, 8）:

- **Normalization:** We employed Root Mean Square Normalization (RMSNorm) in a Pre-Norm configuration, which stabilizes gradients and accelerates convergence compared to standard Layer Normalization. To further prevent attention logits from spiking, we explicitly applied RMSNorm to the Query and Key projections (QK-Norm) before computing attention scores.
- **Positional Encoding:** To better model the translational invariance of biological motifs, we used Rotary Positional Embeddings (RoPE) applied to the query and key vectors at every attention layer.
- **Initialization:** We implemented a Deep Layer Scaling strategy where the weights of residual projection layers were scaled by a factor of $1/\sqrt{2L}$ (where $L$ is the number of layers) (Radford et al., 2019) at initialization. This ensures that the variance of the activations remains bounded as the network depth increases.
- **Training Objective:** The model was trained using a standard autoregressive objective to maximize the log-likelihood of the next token in the discrete protein sequence. The loss function is defined as the Cross-Entropy Loss between the predicted probability distribution over the codebook vocabulary and the ground-truth discrete tokens obtained from the frozen VQ-VAE.
- **Optimization:** We used the AdamW optimizer with β1=0.9, β2=0.95, and an epsilon of 1e-5 to improve numerical stability. The learning rate followed a cosine decay schedule, preceded by a linear warmup for 2,000 steps peaking at 3e-4. To prevent overfitting, we applied a weight decay of 0.01, a dropout rate of 0.1 throughout the network, and gradient clipping at a maximum norm of 0.3. The model was trained for 100M steps on the encoded UniRef50 dataset with an effective batch size of 192.

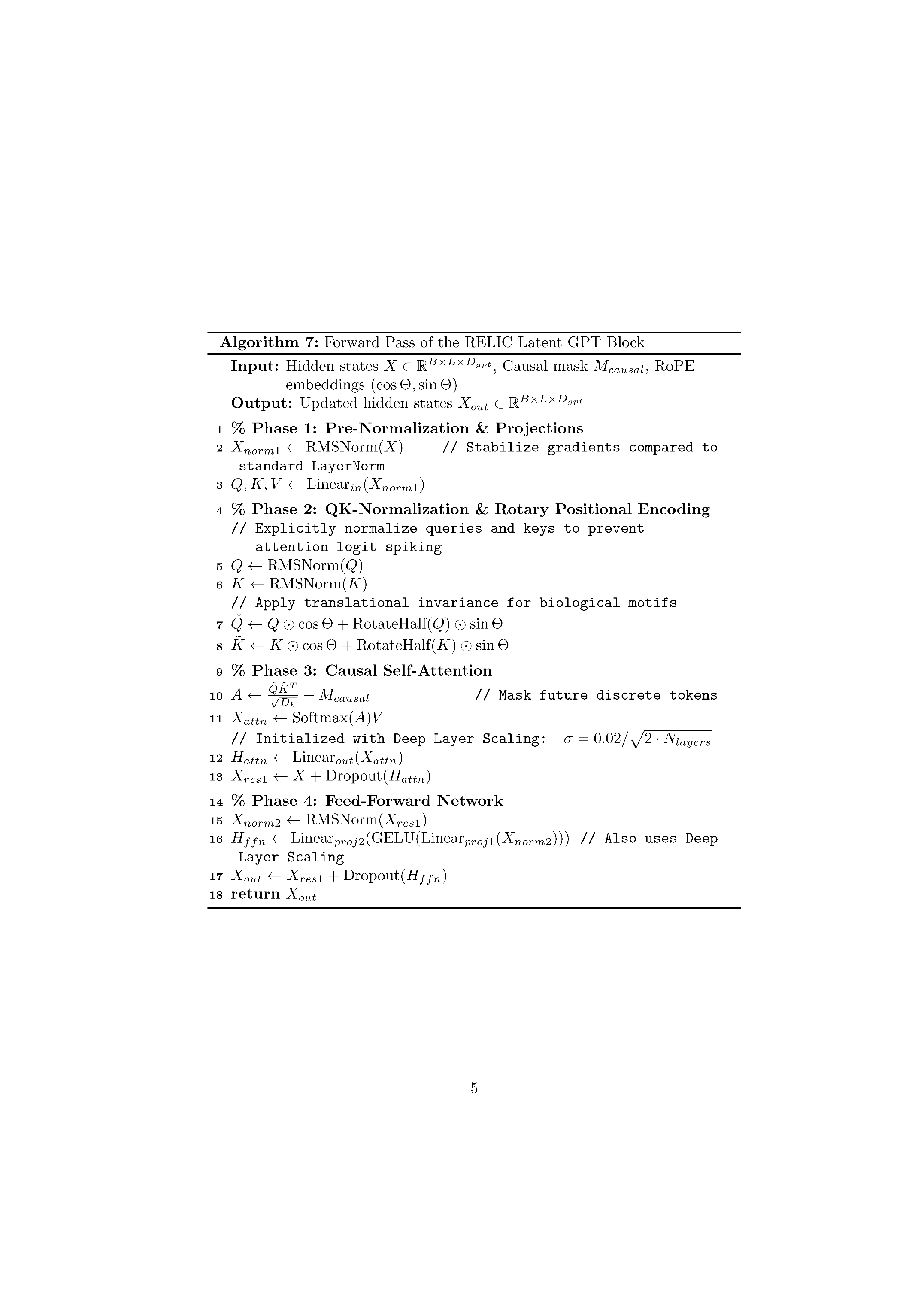

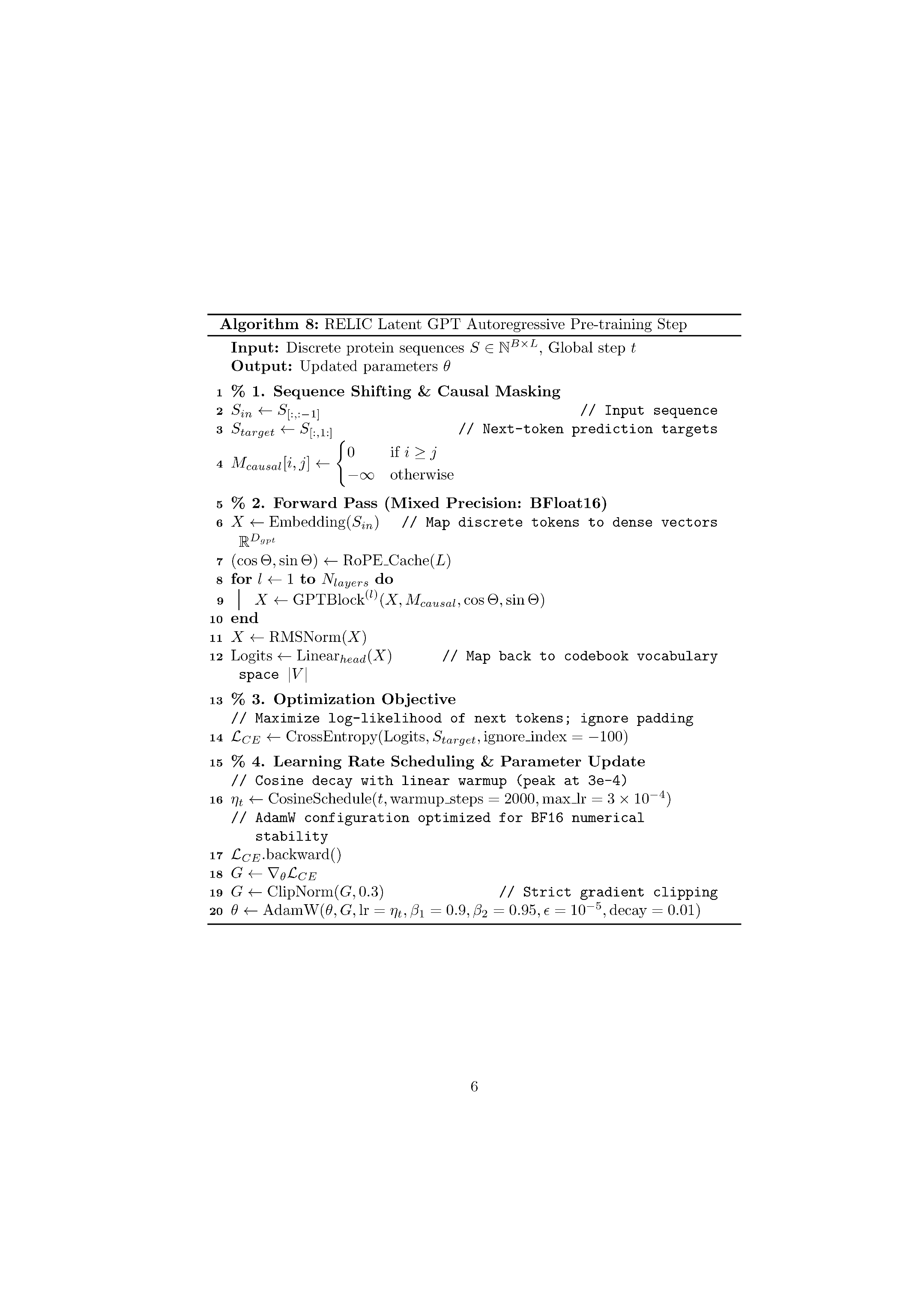

#### Model Usage

##### Latent Space Sequence Alignment

To quantify functional and structural homology between sequences that share low sequence identity, we developed a local alignment algorithm operating directly on the continuous latent representations generated by the model. Unlike standard sequence alignment which relies on discrete amino acid substitution matrices (e.g., BLOSUM62), our method aligns sequences based on the geometric proximity of their post-attention token embeddings.

###### Embedding Whitening and Normalization

Raw embedding spaces from deep learning models are often anisotropic, containing a dominant common signal that obscures specific semantic relationships. To correct for this, we applied a Whitening Transformation to the embeddings.

We computed the global mean vector (μ) and the covariance matrix (Σ) of the token embeddings across a representative subset of the database. Using Singular Value Decomposition (SVD), we derived a whitening matrix $W=US^{-1/2}$, where U ans S are the eigenvectors and eigenvalues of Σ. For a give query embedding sequence $Q\in\mathbb{R}^{L\times D}$ and target sequence $T\in\mathbb{R}^{M\times D}$, the transformed embeddings are calculated as $\hat{x}=(x-\mu)W$. This transformation ensures that the dot product between token vectors reflects their true semantic similarity by isotropically normalizing the latent space.

To efficiently search large-scale databases, we implemented a coarse-to-fine search strategy:

###### Token-Level Coarse Filtering

We utilized a K-Nearest Neighbors (KNN) approach to rapidly identify candidate sequences. For every token in the query sequence, we retrieved the top K = 1000 nearest tokens from the entire database based on cosine similarity. Proteins were ranked by the total count of token-level “hits”, and the top candidates (up to 10,000) were selected for fine-grained alignment. This “bag of tokens” approach effectively filters out unrelated sequences without the computational cost of full alignment.

###### Fine-Grained Latent Smith-Waterman alignment

We performed local alignment on the selected candidates using a modified Smith-Waterman (SW) algorithm (Smith & Waterman, 1981) adapted for continuous vectors. (Algorithm 9)

We defined the similarity matrix $S\in\mathbb{R}^{L\times M}$ as the dot product of the whitened query and target embeddings. To enforce physical realism and reduce noise, we applied Shifter quadratic scoring: If the similarity $S_{i,j}^{'}$ exceeds a threshold $\tau=1.0$, the match score is calculated as ${(S_{i,j}^{'})}^{2}$.

The alignment score $H$ was computed using dynamic programming with affine gap penalties (Gap Open=5.0, Gap Extension=1.0):

$$\left\{ \begin{aligned} 0 \\ H_{i-1,j-1}+Score(S_{i,j}^{'}) \\ H_{i-1,j}-GapPenalty \\ H_{i,j-1}-GapPenalty \end{aligned} \right.$$

To account for length bias, the raw score was first normalized by the geometric mean of the query and target sequence lengths:

$$S_{norm}=\frac{S_{raw}}{\sqrt{L_{q}\cdot L_{t}}}$$

where $L_{q}$ and $L_{t}$ are the full lengths of the query and target sequences, respectively. Finally, to establish statistical significance, we converted the length-normalized scores into Z-scores. We estimated the background score distribution (mean $\mu_{bg}$ and standard deviation $\sigma_{bg}$) by aligning 10,000 randomly selected protein pairs from the database. The final Z-score for a given query-target pair is calculated as:

$$Z=\frac{{Score}_{raw}-\mu_{bg}}{\sigma_{bg}}$$

Alignments were subsequently visualized by tracing the optimal traceback path through the similarity matrix, providing intuitive interpretability for the detected homology.

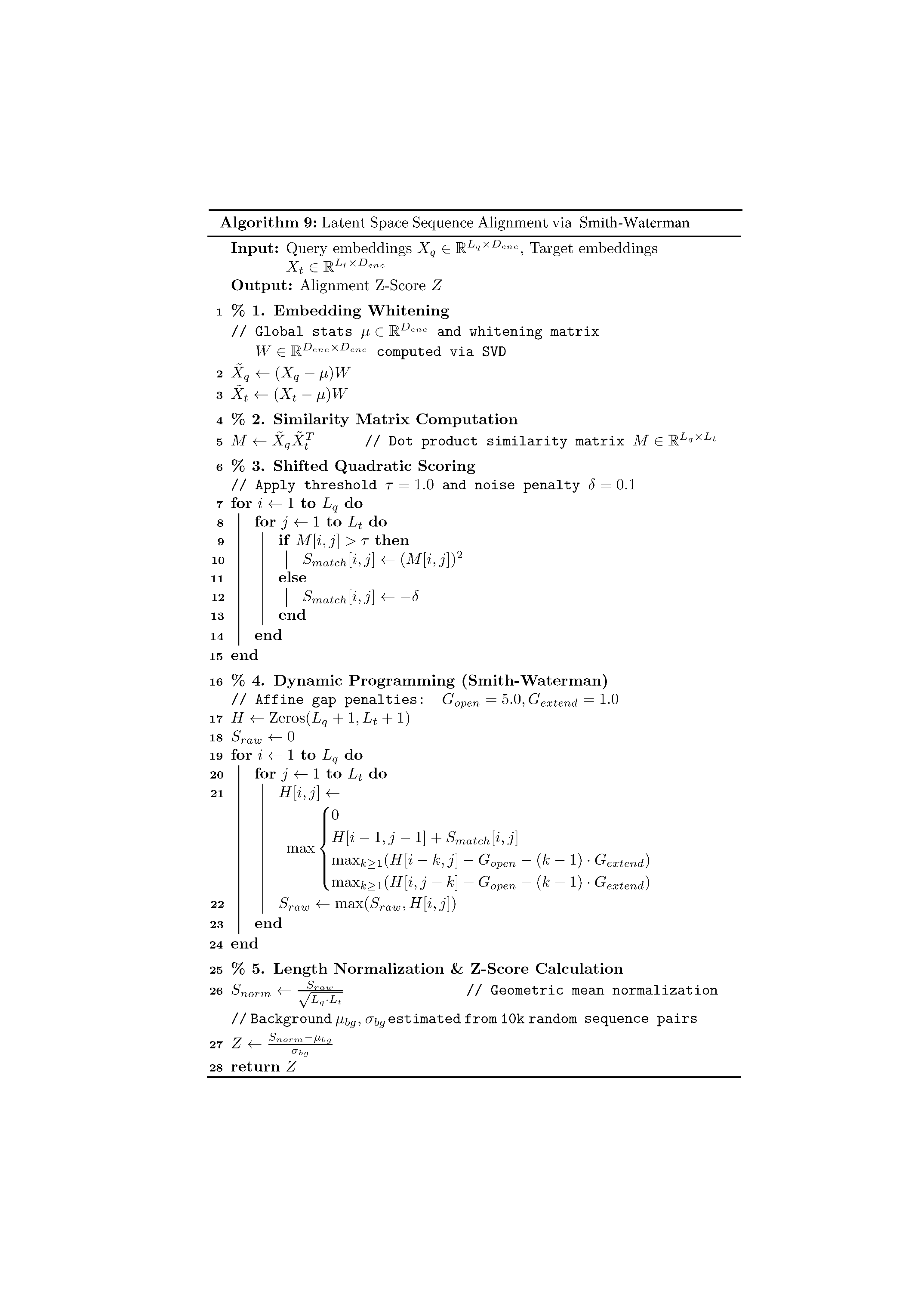

##### Residue level attribution of ProtWords via integrated gradients

To interpret the quantization decisions of the VQ-VAE model and identify amino acids responsible for specific latent protein word assignments, we employed the Integrated Gradients (IG) method (Sundararajan et al., 2017). This method addresses the problem of gradient saturation by accumulating gradients along a linear path from a baseline input to the actual input.

We define the baseline $x^{'}$ as a zero-embedding vector, representing the absence of information. The attribution score for the $i$-th residue, denote as $S_{i}$, is calculated by approximating the path integral using a Riemann sum with $m=50$ steps:

$${IG}_{i}\left( x \right)=(x_{i}-x_{i}^{'})\times\frac{1}{m}\sum_{k=1}^{m} \nabla_{x}\mathcal{L(}x^{'}+\frac{k}{m}(x-x^{'}))$$

Where $x$ is the input embedding and $\mathcal{L}$ is the objective function defined as squared Euclidean distance between the encoder’s continuous latent output $z$ and the target codebook vector $e$:

$$\mathcal{L=}\left\| z-e \right\|_{2}^{2}$$

Finally, to obtain a scalar importance score for each residue, we compute the $L_{2}$ norm of the attribution vector along the embedding dimension. This metric quantifies the sensitivity of the VQ code assignment to specific local features in protein sequence.

##### Case study: Functional annotation using ESMFold2/ESMC SAE features

To benchmark our findings against state-of-the-art continuous protein language models, we queried the uncharacterized C7orf57 (ADMAP1) sequence and C10orf120 (ADMAP2) sequence against the ESMFold2 representations within the recently released ESM Atlas. The prompt and the model's output are reproduced below:

**“Prompt:** Can you predict the function and cellular localization of human protein C7orf57? The sequence of C7orf57 is 'MRNTSKELQGATHRYAPCDWYYHVPVKRSEKAVDAPPASQIPGLSNLGDSHSENLPGTRRYWIKETDSEYVKLAKQGGRPDLLKHFAPGTRKGSPVAYSLPDWYIHHSKPPTASQQEVRAVSMPDYMVHEEFNPDQANGSYASRRGPFDFDMKTVWQREAEELEKEKKKLRLPAIDSKYLSKAGTPLGPKNPAGSRLSFPPVPGQKNSSPTNFSKLISNGYKDEWLQQQQRADSDKRTPKTSRASVLSQSPRDLEGPQDAARLQDAEASEGPEDTPESSQSPEESVSASTPAELK''”

**“Model Output:**

Here's what the ESMC analysis reveals about C7orf57:

**Intrinsic Disorder** — the dominant signal

The low pTM (0.15) paired with a reasonable mean pLDDT (0.73) is a classic hallmark of an intrinsically disordered protein (IDP): local residue-level predictions are plausible, but the model finds no stable global tertiary fold. Every one of the top 10 ESMC SAE features independently converges on the same theme — extended, low-complexity intrinsically disordered regions (IDRs) rich in Ser/Pro/Thr/acidic residues spanning virtually the entire 295-residue sequence.

**Predicted function**

The highest-confidence feature (normalized value 1.05, reliable label) identifies C7orf57 as carrying phosphorylation-rich, acidic C-terminal scaffolding regions that act as multivalent regulatory tails. Reinforcing features at moderate confidence point to:

Phospho-regulated interaction hub: clustered SP/TP dipeptides and phosphosites that recruit modular binders (SH3, WW, 14-3-3 domains)

Scaffold/adaptor role: mediating protein-protein and possibly protein-RNA interactions rather than catalysis

PP1/PKA signaling axis: one feature (value 0.62) suggests the IDR may dock Ser/Thr phosphatases or kinases, consistent with a regulatory subunit-like function

RNA metabolism or chromatin context: multiple features explicitly describe involvement in RNA/RNP complexes or chromatin/replication-checkpoint regulation

Predicted cellular localization

The feature descriptions consistently point toward nuclear and/or nucleocytoplasmic localization — specifically contexts involving chromatin regulation, transcription, RNA metabolism, and replication checkpoints. Some features also mention centrosomal and cytoskeletal scaffolds as alternative contexts, so cytoplasmic localization cannot be fully excluded.

**Caveats**

This protein was folded on-demand (it is not currently an Atlas cluster representative), so no cluster-level comparative data is available. All functional predictions here are inferences from ESMC SAE feature activations, not experimentally validated annotations. The moderate-value features (0.6–0.9) should be treated as hypotheses rather than definitive assignments.

Would you like to search for similar protein clusters in the Atlas to find potential homologs or functionally related proteins? That could help triangulate the biology further. Alternatively, if you have other sequences from your own experiments you'd like to analyze at scale, the ESMC API on biohub.ai supports bulk embedding and SAE feature extraction.”

“**Prompt:**

Can you predict the function and cellular localization of human protein C10orf120? The sequence of C10orf120 is 'MIREWKNDCQRIEKQRASDTMVQERKNEKPVRIFNTNSSFQDQAPTCCQEDLSSASPLRIWSKFYRSDPRIALGKYSPLEKEILRLGGIHTIAARRLLAYKQEEECRMLKELQLLSPDYKQAMEYKKKHSSPCAICVPLEKIWTAKVIAPLEAFKMPQREQVNVSKHIERMRLARALGNHQPLPYIERFTRSSFLSGVGLGPMAKNKARRKEDNYDTHNCDDANQDKKEEAEGKNTKRREIKMNVVFKSKEPKKCLTYHGNDRKSFLPAKKPERSIAGLTNRNLFCISEFPGDLMLMNQDFISRRDHFSDLVKTYSLEEESIWKERMRKATPYHY'”

“**Model Output**:

Here's what the ESMC SAE features and structural data reveal about C10orf120:

**Overall character** — intrinsically disordered protein The structure prediction (pTM 0.207, low; mean pLDDT 0.60, medium) is consistent with a predominantly disordered protein — low pTM reflects the absence of a stable globular fold, not a prediction failure. The top ESMC SAE feature (index 5009, "Polyampholytic low-complexity regions", value 1.1 — high reliability) fires across the entire 335-residue sequence, and three further moderate-reliability IDR features (charged low-complexity IDRs, PTM-dense regulatory IDRs, Pro/Ser-rich disordered scaffolds) also span the full length. This is a strong, convergent signal that most of the sequence is intrinsically disordered.

**Predicted function** — regulatory scaffold Several features point toward a scaffolding/regulatory role:

- "PTM-dense regulatory IDRs" (feature 2639, moderate) highlights SP/TP motifs typical of proline-directed kinase phosphorylation sites, suggesting the protein may be regulated by phosphorylation
- "Pro/Ser-rich disordered scaffolds" (feature 5464, moderate) suggests short linear motifs (SxIP, PxxP, PPxY) that could recruit SH3/EVH1/WW-domain partners
- "Charged helical docking segments" (feature 2440, moderate) hints at partner docking via coiled-coil or amphipathic helical elements

**Predicted cellular localization** — nuclear/chromatin Two moderate-reliability features (2639 and 4155) explicitly describe their associated proteins as nuclear/chromatin or transcriptional regulators. Feature 5009 also calls out "nuclear/cytosolic scaffold and regulatory proteins." The C-terminal tail (residues 304–334) activates a low-reliability feature (7044) for "C-terminal targeting/docking SLiMs" that includes nuclear targeting signals — treat this as a weak hypothesis, not a firm prediction.

**Caveats** UniProt confirms C10orf120 remains uncharacterized, noting only that it is "dispensable for normal development and fertility." The ESMC features converge on a nuclear, IDR-scaffold picture, but no specific binding partners, pathway membership, or enzymatic activity can be inferred from this data alone. The low-reliability features (value < 0.5) should be treated as exploratory hypotheses.

Would you like to search for similar protein clusters in the Atlas to see whether related uncharacterized proteins share these same IDR and nuclear-scaffold signals? That could help contextualize C10orf120 within a broader protein family. If you want to run ESMC embeddings or SAE feature extraction on your own sequence datasets, that's also available via the API on biohub.ai.”

The complete transcript of the ESM Atlas query are available online: C7orf57(<https://biohub.ai/esm/protein/atlas/shared/d66e6283-ee64-4689-b6c8-ec9973b2f5ca>) C10orf120(<https://biohub.ai/esm/protein/atlas/shared/2e9d0e6c-f707-4b34-8337-11766f269117>) and are provided as an HTML document in Supplementary File.

##### ProtWord Distribution Across Species

###### Construction of Species Intensity Vectors

To represent evolutionary characteristics at the species level, we aggregated individual protein encodings into a “vocabulary usage” profile: For each species $S$, we calculated the total frequency $f_{i}$ of each codebook index $i$ across all its constituent proteins. To mitigate the impact of varying proteome sizes and to amplify low-frequency but functionally significant structural motifs, we applied a Log-Frequency transformation:

$$V_{s}=log(1+\sum_{p\in S} count({code}_{i}))$$

To highlight species-specific preferences relative to the background distribution, we normalized the log-counts into Z-scores.

###### Dimensionality Reduction via Independent Component Analysis (ICA)

To isolate independent biological signals from the high-dimensional codebook space, we employed ICA on species-features. The species-feature matrix $X\in\mathbb{R}^{M\times N}$ (where M is the number of species and N=8192) was centered and whitened using FastICA (Hyvarinen, 1999). Unlike Principal Component Analysis (PCA) which maximizes variance, ICA seeks to decompose the data into statistically independent components (ICs) that follow a non Gaussian distribution, better representing distinct evolutionary trajectories. The resulting components were ranked based on the magnitude of their weights in the mixing matrix.

###### Functional Annotation and GO Enrichment Analysis

To elucidate the biological significance of specific code ensembles identified in the evolutionary analysis, we developed a high-throughput functional enrichment pipeline based on the Hypergeometric distribution. We performed a taxonomic audit of the reference database to ensure it reflects a broad evolutionary spectrum. (Fig. S4C).

During the GO enrichment analysis, all sequences containing at least one code from the target ensemble are aggregated into the sample subset of size $N$. For each GO term, we assess its occurrence in the sample subset ($k$) relative to its occurrence in the total background ($n$) of size $M$. The statistical significance is calculated as:

$$P\left( X\geq k \right)=\sum_{i=k}^{min(n,N)} \frac{(\begin{matrix} n \\ i \end{matrix})(\begin{matrix} M-n \\ N-i \end{matrix})}{(\begin{matrix} M \\ N \end{matrix})}$$

This test determines whether a structural motif is associated with a specific biological function more frequently than expected by random chance. To control the Type I error rate inherent in testing thousands of GO terms, we applied Bonferroni correction.

###### Prediction and Analysis of Intrinsically Disordered Regions (IDRs)

To characterize the relationship between protein words and intrinsic disorder, we predicted per-residue disorder probabilities using Metapredict V2 (Emenecker et al., 2021). For each protein sequence in the dataset, Metapredict generated a vector of disorder scores. Protein words were mapped to protein sequences using a downsampling scale of 4 residues per word. For every occurrence of a specific word at index k in the word sequence, the corresponding physical center in the protein sequence was defined as $C_{k}=4k+2.$

To visualize the disorder landscape associated with each code, we extracted local disorder profiles using a window of radius R=30 residues. For each word type c within a species, we aggregated all corresponding windows to calculate the position-wise mean and standard deviation of IDR probabilities:

$$\mu_{j}=\frac{1}{N_{c}}\sum_{n=1}^{N_{c}} s_{n,j} and \sigma_{j}=\sqrt{\frac{1}{N_{c}}\sum_{n=1}^{N_{c}} {(s_{n,j}-\mu_{j})}^{2}}$$

Where $N_{c}$ is the total count of code c, and j represents the relative position within the window ($j\in[-30,+30]$).

To systematically identify codes significantly enriched in disordered regions, we defined a ‘local disorder score’ for each code occurrence as the mean IDR probability within a narrow core window ($C_{k}\pm2$).

###### Pfam Annotation

Protein domains were annotated using the HMMER3 suite (v3.4) (Johnson et al., 2010) against the Pfam-A database (release 35.0). We performed proteome-wide searches using the hmmsearch command. To ensure high-confidence assignments, hits were filtered based on the domain independent E-value (Evalue < ${10}^{-5}$). In cases of overlapping domain predictions, the conflict was resolved by retaining the hit with the lowest E-value, ensuring a non-redundant domain architecture for each protein.

##### Autoregressive Protein Sequence Generation with Latent Semantic Guidance

To generate novel protein sequences, we employed an autoregressive decoding strategy using the pre-trained latent GPT. The generation process was initiated with a specific start-of-sequence (<sos>) token. At each decoding step t, the model predicted the probability distribution of the next latent token based on the preceding context. To balance sequence diversity and biological plausibility, we implemented a composite sampling strategy. Specifically, the logits were scaled by a temperature factor T, followed by Top-k filtering and Nucleus sampling (Top-p) to truncate the low-probability tail of the distribution.

To mitigate the generation of chemically incoherent motifs, we introduced a novel latent semantic guidance mechanism. Utilizing the pre-trained VQ-VAE codebook embeddings, we enforced a geometric constraint in the latent space: candidate tokens were filtered to maintain a cosine similarity greater than a threshold with the token possessing the highest predicted probability. This ensures that sampled tokens remain within a semantic neighborhood of the model’s most confident prediction. Furthermore, a repetition penalty was applied to prevent degenerative repetitive patterns. The generated latent indices were subsequently mapped back to the amino acid space using the VQ-VAE decoder. (Algorithm 10)

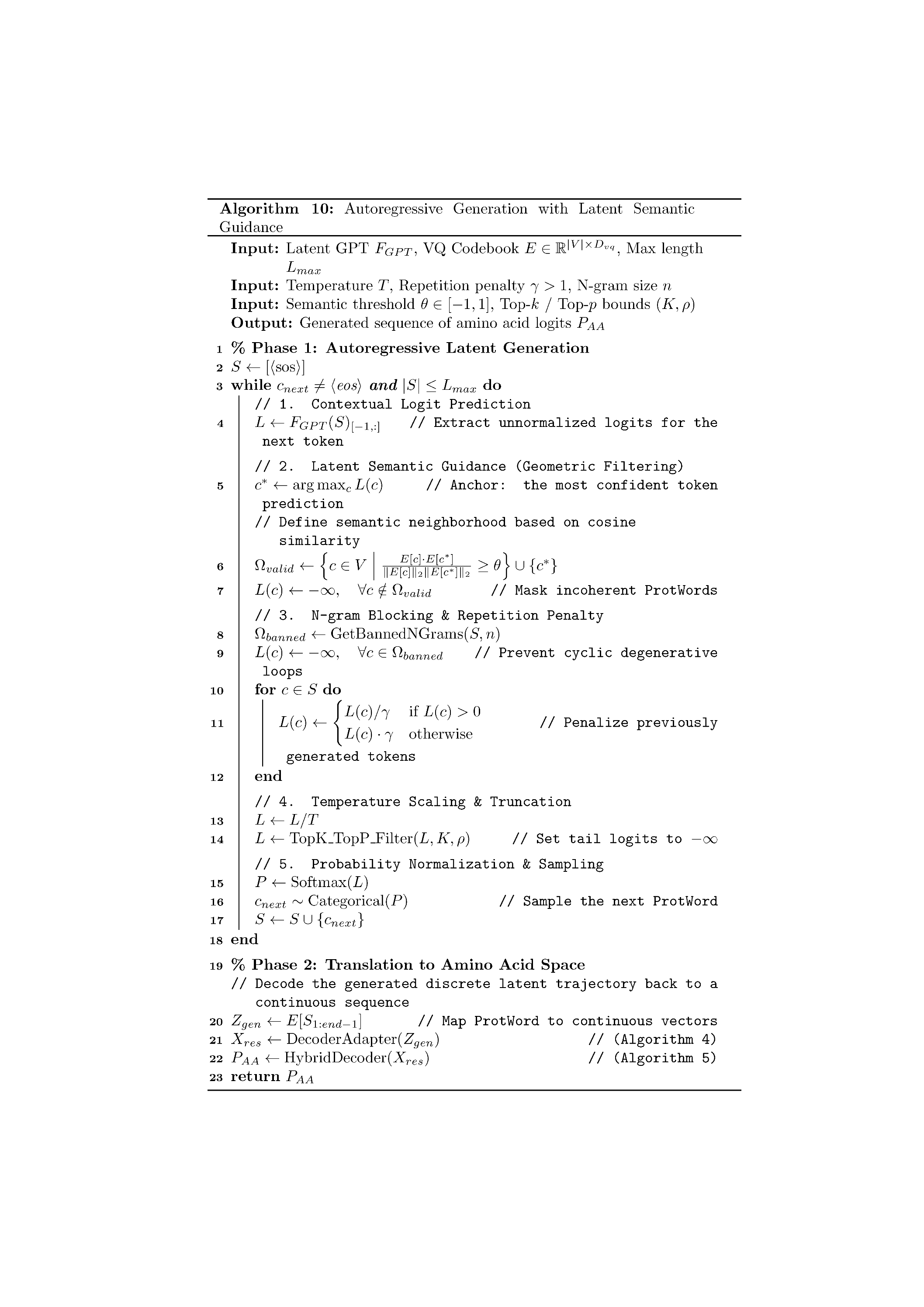

##### Fine-tuning with Cofilin Family Sequences

To enable the targeted generation of specific functional families, we fine-tuned the pre-trained ProteinGPT model on curated datasets of and Cofilin/ADF homologs.

- **Model Adaptation:** The fine-tuning process was initialized from the pre-trained general-purpose checkpoint. Since the pre-training task was unconditional, no architectural modification was required, and we directly resumed autoregressive training on the family-specific subsets.
- **Hyperparameters and Training Dynamics:** To avoid catastrophic forgetting while adapting to the new sequence distribution, we employed a lower learning rate and a smaller batch size compared to the pre-training phase. Fine-tuning was performed using the AdamW optimizer with a peak learning rate of 1e-5 and a weight decay of 0.01. We utilized a cosine learning rate decay schedule with a short linear warmup period (10% of total steps) to stabilize the early phase of domain adaptation.
- **Regularization and Optimization:** To prevent overfitting on the smaller fine-tuning dataset, a network-wide dropout rate of 0.1 was maintained. To ensure numerical stability, gradients were strictly clipped at a maximum L2 norm of 1.0, and all forward passes were executed using BFloat16 mixed precision. The model was trained with a global batch size of 96 sequences to maintain stable gradient estimation, and the fine-tuning process proceeded for a total of 450 steps.

##### Filtering and Benchmarking of Designed Proteins

To evaluate the structural novelty and foldability of the generated proteins, we implemented a dual-stage filtering strategy based on sequence redundancy and structural prediction confidence.

- **Sequence Redundancy Filtering:** To ensure the design novelty, we performed all-versus-all sequence alignment against the UniRef50 database (Release 2025_04) using MMseqs2. We applied a filtering criterion to retain sequences with a global sequence identity ranging between 0.3 and 0.7 relative to existing natural sequences. This range excludes both trivial memorization (identity > 0.7) and non-functional, excessively divergent designs (identity < 0.3).
- **Structural Foldability Assessment:** We utilized ESMFold (Lin et al., 2023) to predict the structural models of both unconditional designed proteins and pwCofilin variants. Foldability was benchmarked using the predicted TM-score (pTM) and the mean predicted local distance difference test (pLDDT). Sequences exhibiting a pTM score above 0.7 were selected for subsequent experimental validation.
- **Structural Visualization and Modeling:** Structural models presented in the figures were either obtained from the AlphaFold Database (<https://alphafold.com/>) (Fleming et al., 2025) or generated using the AlphaFold3 server (<https://alphafoldserver.com/>) (Abramson et al., 2024). For proteins involving metal-ligand coordination, such as Fe–S cluster binding proteins, structures were predicted using a local deployment of AlphaFold3, explicitly incorporating the ligand via Chemical Component Dictionary (CCD) representation.
- **Homology Assignment:** To accurately identify the nearest natural neighbors for the pwCofilin-7, -14, and -90 variants, we performed BLASTp searches against the UniRef100 database. Among the retrieved hits, we selected the entry with the lowest E-value to determine the sequence identity. Pairwise sequence alignments were generated using the UniProt alignment tool, which employs the Clustal Omega tool (Sievers et al., 2011) with default parameters. Multiple sequence alignments (MSAs) were visualized using ESPript 3.2 (Gouet et al., 1999), incorporating the secondary structure annotation derived from the structure of human Cofilin-1 to map functional residues onto the conserved protein fold. The F/G binding site, were identified based on the structural mapping described by (Kraus et al., 2022).

##### Phylogenetic Analysis

To determine the evolutionary relationships between the functional designs and the natural protein landscape, we reconstructed a maximum likelihood (ML) phylogenetic tree.

First, to mitigate computational redundancy, we clustered the combined training and validation sets using MMseqs2 (similarity threshold 0.5, coverage 0.8). These representative natural sequences were then aligned with our functional designs using MAFFT (v7.525) (Katoh et al., 2002). We utilized the --auto strategy, which dynamically selected the FFT-NS-2 algorithm and performed two iterations of iterative refinement to optimize the alignment accuracy for the 324-sequence dataset.

Phylogenetic reconstruction was conducted using IQ-TREE (v3.1.2) (Nguyen et al., 2015). We employed the integrated ModelFinder tool to objectively determine the best-fit substitution model based on the Bayesian Information Criterion (BIC). The model Q.PFAM+R6 was identified as optimal, incorporating a FreeRate model with 6 rate categories to accurately account for among-site rate heterogeneity. The ML tree was constructed using IQ-TREE's standard approach, and the final phylogeny was visualized and annotated using iTOL (<https://itol.embl.de/>) (Letunic & Bork, 2021).

#### Cellular and in vitro Experiments

##### Plasmid Construction

The cDNAs encoding ADMAP1, ADMAP2, and human CFL1 (NCBI Reference Sequences: NM_001100159.3, NM_001010912.4, and NM_005507.3, respectively), along with the engineered cofilin variants, were synthesized de novo (BGI Genomics). For mammalian cell transfection assays, the ADMAP1 and ADMAP2 sequences were fused to an EGFP-Flag tag, whereas the human CFL1 and engineered cofilin sequences were fused to a single Flag tag; all constructs were subsequently cloned into the pcDNA3.1 expression vector. For recombinant protein purification, these tagged sequences were subcloned into the pET21 backbone by homology recombination (HR) using In-Fusion HD Cloning Kit (Takara Bio, #639650). For All plasmids were isolated and purified using the Qiagen Plasmid Miniprep Kit (Qiagen, #12123).

##### Cell Culture, Transfection, and Drug Treatments

HeLa cells were cultured in DMEM (ThermoFisher, #C11995500BT) and mIMCD3 cells in specialized mIMCD3 medium (Immocell, #26012619), both supplemented with 10% FBS (Gibco, #B210750RP) and 1% penicillin-streptomycin (Gibco, #268394) at 37°C with 5% CO₂.

For HeLa cell imaging, cells were transfected with a plasmid encoding the EGFP-fused predicted human microtubule-binding protein using Lipofectamine 3000 Transfection Kit (ThermoFisher, #3241883). For mIMCD3 ciliary assays, cells were co-transfected with 250 ng of ARL13B-mCherry (ciliary marker) and 250 ngEGFP-fused predicted human microtubule protein plasmid. Six hours post-transfection, serum starvation was induced by replacing the growth medium with DMEM/F12 (1:1) medium (Yuanye, #KR404921) containing 5% FBS (Gibco, #B210750RP) for 24 h to stimulate ciliogenesis prior to fixation. For cofilin localization assays, cells were transfected with a plasmid expressing Flag-tagged engineered cofilin. All samples were processed for fluorescence observation 24 h post-transfection or post-treatment.For live-cell visualization of the microtubule network, HeLa cells expressing ADMAP1-EGFP or ADMAP2-EGFP were incubated with Tubulin Tracker Deep Red (ThermoFisher, #T34077, 1:1000) for 30 min at 37°C with 5% CO₂. Following incubation, the cells were gently washed with pre-warmed PBS prior to live-cell imaging.

##### Immunofluorescence

Following specific treatments, the culture medium was aspirated, and cells were fixed with 4% paraformaldehyde (PFA;Yuanye, #R20497) for 15 min at room temperature. After three brief washes with PBS, cells were permeabilized with permeabilization buffer (Beyotime, #P0096) for 10 min and incubated in blocking buffer (Beyotime, #P0260) for 1 h at room temperature.

To determine the localization of engineered cofilin, cells were incubated overnight at 4°C on a laboratory shaker with an anti-Flag primary antibody (Yeasen, #30504ES50) diluted in blocking buffer. Samples were subsequently washed three times with PBS (5 min per wash) under agitation. Cells were then incubated with secondary antibodies (Dylight 649, Goat Anti-Rabbit IgG; 1:300; Abbkine, #A23620) and iFlour 555 phalloidin (1:500; Yeasen, #40737ES75) in a dedicated antibody dilution buffer (Beyotime, #P0265) for 1 h at room temperature in the dark. After three final washes with PBS under agitation, coverslips were prepared for imaging.

##### Fluorescence Imaging and Analysis

Super-resolution and confocal images were acquired 24 h post-transfection or post-treatment using a Zeiss LSM 900 confocal microscope equipped with an Airyscan detector (Zeiss) and a 63×/1.40 oil-immersion objective . The system was operated via ZEN software (Zeiss). EGFP signal was excited using a 488 nm laser line, while mCherry, rhodamine, and phalloidin signal were excited via a 561 nm laser line. Additionally, the Tubulin Tracker Deep Red (ThermoFisher, #T34077) signal was excited using a 640 nm laser line. This setup was utilized to sequentially image protein subcellular localization, ciliary morphology in mIMCD3 cells, and the actin cytoskeleton.

For morphological quantification, a minimum of 200 transfection-positive cells per condition were randomly selected across multiple independent fields of view. The percentage of cells exhibiting abnormal actin cytoskeletal structures was subsequently determined based on phalloidin staining patterns.

##### Preparation of Cell Lysates

HeLa cells were cultured in two 10-cm dishes and transfected with 10 μg of ADMAP1-EGFP-Flag plasmid per dish. At 24 h post-transfection, cells were washed with pre-warmed PBS, detached using 0.25% trypsin (Gibco, #3203949), and harvested via centrifugation at 453 × g for 10 min at 4°C. To prepare the cellular extracts, cell pellets were resuspended in 150 μL of ice-cold lysis buffer comprising BRB80 (80 mM K-PIPES pH 6.8, 1 mM EGTA, 1 mM MgCl₂), 0.1% Triton X-100 (Sigma-Aldrich, #T9284), and a protease inhibitor cocktail (Roche, #5892791001). Mechanical cell disruption was strictly performed on ice by continuous trituration using an insulin syringe (KDL, #u100) for 10 min to induce shear stress. The resulting homogenates were subjected to ultracentrifugation at 33,800 × g for 25 min at 4°C using an ultracentrifuge (Beckman Coulter, #MAX-XP) equipped with a TLA-55 rotor (Beckman Coulter). The clarified supernatants containing the target ADMAP1-EGFP-Flag proteins were aliquoted, snap-frozen in liquid nitrogen, and stored at −80°C until further biochemical evaluation.

##### In Vitro Microtubule Assay

Purified ADMAP1 was mixed with rhodamine-labeled tubulin in a 6.25-μL reaction containing 1× BRB80 buffer (80 mM PIPES-K pH 6.8, 1 mM MgCl₂, 1 mM EGTA), 1 mM GTP (Roche, #10106399001), an additional 4 mM MgCl₂, and 4% anhydrous DMSO (Solarbio, #ID9010). The mixture was incubated on ice for 5 min, followed by polymerization at 37°C for 30 min in a thermal cycler. Assembled microtubules were stabilized by adding 250 μL of 1× BRB80 containing 20 μM taxol (Sigma-Aldrich, #T7191) and pelleted via centrifugation at 14,000 rpm for 20 min at room temperature. The supernatant was removed, and the pellet was resuspended in 200 μL of 1× BRB80 with 20 μM taxol for imaging.

##### Purification of recombinant ADMAP1 and EB3

For expression and purification of recombinant C7orf57-EGFP-flag and EB3-mCherry-StrepII, plasmids were transformed into BL21 (DE3) bacteria. 10-mL culture from a single colony was used to inoculate a 1-L fresh LB medium for further incubation at 37 °C with shaking. After 3.5-4 h, IPTG was added to the culture to a final concentration of 1 mM to induce protein expression at 16 °C for 16 h. Then, bacteria pellets were collected by centrifugation and resuspended in 50 mL of ice-cold lysis buffer (80 mM PIPES, 100 mM KOH, 500 mM NaCl, 1 mM EGTA, 1mM MgCl_2_, pH 6.9) containing 1 mM PMSF. After breaking bacteria by microfluidizer (AH-TITAN PLUS) at 800 bar for 10 min, the lysate was centrifuged at 12,000 rpm for 30 min at 4 °C. The supernatant was transferred to a fresh tube containing 0.5 mL of 50 % slurry of Anti-DYKDDDDK (for C7orf57) or Streptactin (for EB3) agarose beads (Smart-Lifesciences). After rotating at 4 °C for 2 h, the mixture was transferred to a column to allow the settlement of the beads. The column was washed with 20 volumes of lysis buffer, and eluted with 5 volumes of elution buffer [lysis buffer with 0.25 mg/mL 3xFlag peptide (for C7orf57) or 2.5 mM D-biotin (for EB3)]. Eluates were concentrated to 50 μL with Amicon Ultra 30K device (Millipore) at 4 °C. 3xFlag peptide or D-biotin was removed through buffer exchange during the ultrafiltration process. BCA assay was used to measure the final concentration of proteins. The proteins were divided into 5 μL aliquots, snap-frozen in liquid nitrogen and stored at -80 °C.

##### In vitro microtubule (MT) dynamics assay

In vitro MT dynamics assay followed the protocol described previously (Zhang et al., 2022). Briefly, the flow chamber was incubated with 0.2 mg/mL PLL-PEG-biotin (Susos AG) and then with 1 mg/mL neutravidin (Invitrogen) in MRB80 buffer (80 mM PIPES, 100 mM KOH, 1 mM EGTA, 4mM MgCl_2_, pH 6.9). Subsequently, Hilyte647-conjugated and biotin-conjugated MT seeds were attached to the coverslip through biotin-neutravidin interactions. The flow chamber was blocked with 1 mg/mL κ-casein. The reaction mixture, which consisted of MRB80 buffer containing C7orf57-EGFP-flag (1 μM), EB3-mCherry (50 nM), porcine brain tubulins (20 μM) (Cytoskeleton), 2 mM GTP, 0.2 mg/ml κ-casein, 2% PEG-8,000, and oxygen scavenger mix (50 mM glucose, 400 μg/ml glucose oxidase, 200 μg/ml catalase, and 4 mM DTT) was added to the chamber. The flow chamber was sealed with vacuum grease and imaged immediately at 30 °C using a Nikon Eclipse Ti-E inverted microscopy equipped with a 100×/1.4 objective. Images were acquired at the TIRF (total internal reflection fluorescence) mode with reflection angle of 65°, laser wavelength of 488 nm / 561 nm / 640 nm and 8 sec intervals.

#### Animal Experiments

##### Generation of Knock-out (KO) Mouse Lines

All procedures for the care and use of laboratory animals was approved by the Institutional Animal Care and Use Committee (IACUC) and Internal Review Board of Tsinghua University (Animal Protocol number: 24-OGS2). C7orf57 (+/-) mice were generated from wild-type C57BL/6 animals using CRISPR-Cas9 by pronuclear injection. The microinjection mix consisted of 20 ng/μL guide RNA and 20 ng/μL Cas9 mRNA. Founders were genotyped by Sanger sequencing of genomic DNA extracted from mousetail tissues, using C7orf57-specific primers. Founder mice (+/-) were then crossed with C57BL/6 mice, and the heterozygous offsprings were intercrossed to produce homozygous KO lines, which were confirmed by Sanger sequencing. Animals were maintained in SPF environment. The guide DNA sequences used for microinjection: ATCCAGTACCTCCGAGCACG.

##### Immunofluorescence

For IF staining, mouse sperms in PBS buffer were spreaded on poly-L-lysine -coated glasses. After incubation for 10 min at room temperature, the sperms were fixed by 4 % paraformaldehyde (PFA) for 10 min, permeabilized by 0.1 % Triton X-100 for 10 min, and then blocked with QuickBlock™ Blocking Buffer (Beyotime) for 1 hour. Blocking buffer containing anti-C7orf57 1st antibody (Abcepta, #AP55914) (1:200 diluted) was incubated with sperms overnight at 4 ℃. After washing with PBS for 3 times, Secondary Antibody Dilution Buffer (Beyotime) containing Dylight488-conjugated goat-anti-rabbit 2nd antibodies (1:300 diluted) were incubated with sperms for 1 hour at room temperature, followed by washing 3 times with PBS buffer and staining with 5 μg/mL Hoechst33342. Imaging was performed using an Axio Observer Z1 microscope (Carl Zeiss) equipped with 488 and 405 laser lines, a Yokogawa spinning disk head, an Andor iXon + EM-CCD camera, and a Zeiss 100×/1.46 objective.

##### Sperm Isolation

Male wild-type and GM11992 knockout mice (10 to 13 weeks of age; n ≧ 3 per genotype) were euthanized via cervical dislocation. Following abdominal sanitization with 70% ethanol, bilateral epididymides were excised and immediately transferred into 3.5-cm dishes (ThermoFisher, #150460) containing pre-warmed PBS.

To facilitate sperm release, the epididymides were minced and incubated in a 37°C water bath for 5 min. Following gentle agitation to ensure a homogenous suspension, 10 μL of the sperm suspension was loaded into a disposable sperm counting slide (Mailang). Sperm motility was immediately observed and recorded via video using a microscope equipped with a 10× objective. Video acquisition was performed using Suijia software (**Suijia**).

##### Sperm Motility Analysis and Particle Tracking

Sperm motility was quantitatively analyzed using custom Python scripts based on the OpenCV and Trackpy libraries. Raw video files were first converted to grayscale, and a static zero-thresholding method (threshold intensity = 80) was applied to reduce background noise and isolate sperm heads.

To reconstruct sperm trajectories, particle identification and linking were performed using Trackpy. Sperm heads were localized in each frame by detecting features with a defined apparent diameter of 17 pixels and a minimum integrated intensity (mass) of 1000. Subsequently, identified features were linked across consecutive frames to form continuous trajectories. The linking algorithm was configured with a maximum search range of 20 pixels and a memory parameter of 25 frames to account for temporary particle occlusion or transient detection failures. Adaptive search parameters (adaptive step = 0.95, adaptive stop = 5) were utilized to optimize tracking accuracy. Short, non-informative trajectories spanning less than 100 frames were discarded from further analysis.

Kinematic parameters were calculated for each validated trajectory by converting pixel coordinates to physical dimensions. The Curvilinear Velocity (VCL) was determined by calculating the instantaneous frame-to-frame speed, which was then smoothed using a centered rolling average with a 5-frame window. The Straight-Line Velocity (VSL) was calculated as the Euclidean distance between the initial and final position of a sperm track divided by the total trajectory duration. The Linearity (LIN) of the movement was defined as the ratio of mean VSL to mean VCL.

To ensure the accuracy of active motility measurements, a stringent computational filter was applied to eliminate artifacts caused by passive fluid drift. Trajectories were classified as drift and excluded from the final dataset if they simultaneously exhibited high directionality (trajectory angle between -70° and -110°), high linearity (LIN > 0.7), and low average velocity (mean VCL < 60 $\mu m/s$). Finally, statistical comparisons of the kinematic parameters across different experimental groups were performed using non-parametric statistical tests.

##### TEM of Mouse Sperm.

The samples were diluted in 0.1M phosphate buffer (pH 7.4), and spermatozoa were collected by centrifugation. The pellets were loaded onto 100-μm-deep specimen carriers using a pipette, briefly air-dried, and covered with sapphire discs. This assembled “sandwich” was mounted in the standard 3 mm sample holder and rapidly frozen with an HPM100 high-pressure freezing system (Leica Microsystems). Frozen samples were subsequently transferred into 1.5 mL cryovials containing 1 mL of a solution of 1% (w/v) osmium tetroxide and 0.1% (w/v) uranyl acetate (Electron Microscopy Sciences) in acetone. Freeze substitution was carried out in an AFS2 machine (Leica Microsystems) using the following program: -90°C for 72 hours, -60°C for 24 hours, -30°C for 18 hours, followed by a gradual increase to 4°C over 4 hours.

After substitution, the samples were washed three times with pre-cooled pure acetone and then brought to room temperature. Resin infiltration was performed using SPI-Pon 812 resin (Structure Probe Inc.) with sequential steps: 50% resin (in acetone) for 2 hours, 75% resin overnight, 100% resin for 2 hours, fresh 100% resin for 6 hours, and a final change of 100% resin for 1 hour. Samples were then embedded in flat molds, and polymerized at 60°C for 3 days.

Ultrathin sections (90 nm) were obtained using a UC7 ultramicrotome (Leica Microsystems) and picked onto 200-mesh copper grids (Gilder Grids Ltd.). Sections were post-stained with 2% uranyl acetate and Reynolds’ lead citrate, and imaged on an FEI Tecnai G2 Spirit transmission electron microscope operated at 120 kV (Thermo Fisher Scientific Co.). Definitions and classification criteria for sperm phenotypes were adopted according to (Chemes et al., 1998; Mortimer, 2018).

#### Biochemistry Experiments on Generated Cofilin Activity

##### Plasmid Construction

For prokaryotic expression, gene fragments encoding *Escherichia coli* Cofilin isoforms (C7, C14, and C90) and human Cofilin-1 (S3A mutant) were synthesized following codon optimization. These sequences were subsequently subcloned into the pET.M.3 expression vector, engineered to introduce a C-terminal Strep-tag fusion to facilitate subsequent protein purification.

##### Bacterial Protein Expression and Purification

The recombinant plasmids were transformed into *E. coli* BL21(DE3) competent cells for protein production. Protein expression was induced by adding isopropyl β-D-1-thiogalactopyranoside (IPTG) to a final concentration of 0.5 mM, followed by incubation at 16 °C overnight. Harvested bacterial cells were resuspended and lysed in ice-cold lysis buffer containing 40 mM Tris-HCl (pH 8.0), 250 mM NaCl, 5% glycerol, 1 mM dithiothreitol (DTT), and a complete protease inhibitor cocktail (Roche). The clarified lysate was subjected to affinity purification using Strep-Tactin Sepharose beads. Following incubation, the resin was washed extensively with wash buffer (40 mM Tris-HCl, pH 8.0, 250 mM NaCl, 5% glycerol, 1 mM DTT). Bound proteins were eluted with elution buffer supplemented with 2.5 mM desthiobiotin. Protein concentrations were determined by comparing sample band intensities against a bovine serum albumin (BSA) standard curve via SDS-PAGE. Gels were stained with Coomassie Brilliant Blue, imaged using a CCD camera, and quantitative densitometry was performed using ImageJ software.

##### F-Actin co-sedimentation assay

Monomeric G-actin was polymerized into filamentous actin (F-actin) prior to the severing assay. Briefly, G-actin was converted to Mg-actin by the addition of 10× ME buffer (yielding final 1× concentrations of 1 mM EGTA and 0.1 mM MgCl₂) and incubated for 5 min at room temperature. Polymerization was subsequently induced by adding 10× KMEI buffer (final 1× concentration: 50 mM KCl, 1 mM MgCl₂, 1 mM EGTA, 10 mM imidazole, pH 7.0) and 1× Buffer G (5 mM Tris-HCl pH 8.0, 0.2 mM CaCl₂, 0.2 mM ATP, 0.2 mM DTT, 0.02% NaN₃), followed by a 10-min incubation at room temperature.

To evaluate F-actin severing activity, 1 μM of pre-polymerized F-actin was incubated with 0.5 μM of purified wild-type CFL1 or engineered cofilin variants. The 100 μL reaction was carried out in 1× Buffer G (maintained with 0.2 mM ATP and 0.2 mM DTT) and incubated at 30°C for 1 h.

Post-incubation, the mixtures were subjected to ultracentrifugation at 100,000 × g for 30 min at 22°C using a tabletop ultracentrifuge (Beckman Coulter, #MAX-XP) equipped with a TLA-55 rotor (Beckman Coulter). For each sample, 80 μL of the supernatant fraction was carefully collected and mixed with 5× SDS loading buffer (Yeasen, #20315ES). After discarding the remaining liquid, the pellet fraction was directly resuspended in 1× SDS loading buffer. Both fractions were denatured at 95°C for 5 min, resolved on FastPAGETM SDS-PAGE gels (Tsingke Biotech, #TSP024), and stained with Safe Protein Fast Staining Solution (Tsingke Biotech, #TSP8112) for 10 min. Gels were subsequently washed in PBS for 1 h prior to imaging.

##### In Vitro Actin Dynamics Assays

The filament-severing activity of Cofilin isoforms was assessed using time-lapse total internal reflection fluorescence microscopy (TIRFM), based on established protocols(Tian et al., 2015). Briefly, flow chambers were coated with 25 nM N-ethylmaleimide-myosin to immobilize actin filaments. Subsequently, 140 nM pre-assembled actin filaments (90% rhodamine-labeled) were introduced into the chamber and incubated in the dark for 10 minutes.The flow cell was subsequently washed with 1×TIRFM buffer (10 mM imidazole, pH 7.0, 50 mM KCl, 1 mM MgCl2, 1 mM EGTA, 5 mM DTT, 0.2 mM ATP, 50 μM CaCl2, 15 mM glucose, 20 μg/mL catalase, 100 μg/mL glucose oxidase,0.1% BSA, and 0.5% methylcellulose).To initiate filament severing, the chamber was perfused with 1×TIRFM buffer containing each 100 nM Cofilin.We quantified two characteristic parameters of Cofilin activity: the average severing frequency and the subunit off-rate. Severing frequency was defined as the number of breaks per unit filament length per unit time (breaks/μm/s), as previously described(Andrianantoandro & Pollard, 2006). For analysis, more than 20 actin filaments exceeding 3 μm in length were selected for the quantification. The average subunit off-rate (subunits/s) was calculated by monitoring the shrinkage rate at filament ends, assuming a conversion factor of 334 subunits per micrometer of filament length. The pointed and barbed ends of the actin filaments were distinguished according to prior reports (Shi et al., 2013).
